# Nutrient availability dictates cancer metabolism-based therapeutic responses of non-oncology drugs

**DOI:** 10.1101/2025.01.05.631340

**Authors:** Woo Yang Pyun, Jae Hyung Park, Jae Won Roh, Dongkyu Jeon, Jongwan Kim, Ji Eun Paik, Seok Chan Cho, So Yeon Park, Hocheol Lim, Hyungwoo Kim, Young Jin Jang, Jaehoon Lee, Soo-Youl Kim, Kun-Liang Guan, Heon Yung Gee, Han-Woong Lee, Kyoung Tai No, Han-Sol Jeong, Wan Namkung, Joo Hyun Nam, Hyun Woo Park

## Abstract

Metabolic deregulation is a major hallmark of cancer, therefore, interventions that modify tumor nutrient availability are considered attractive adjuvants for improving clinical outcomes for cancer patients. Much work remains, however, in clarifying how the nutritional status of each patient can affect the metabolic vulnerability of drugs and inform individual medication guidelines. Working toward the goal of oncometabolic precision medicine, we introduce CM-SLP (cancer metabolism-based synthetic lethality platform), a high-throughput screening platform that explores the metabolic vulnerability of non-oncology drugs induced by altered nutrient availability and predicts the potential synthetic lethal interactions with either hyper- or hypo-nutrient conditions. We present promising CM-SLP candidates, such as propafenone and biguanides, as representative non-oncology drugs that cooperatively enhance cytotoxicity via dysregulated metabolic pathways. Furthermore, identifying mTOR and Hippo pathways as mediators of combined propafenone/hypoglycemia or biguanides/hypoglycemia treatments, respectively, we were able to circumvent the need for dietary interventions by administering the mTOR or TEAD inhibitors to induce energy stress and cancer cell death. Together, CM-SLP represents a critical step toward integrating metabolic profiling into precision oncology, offering novel therapeutic avenues tailored to individual patient needs.

## Introduction

Cancer progression depends on reprogrammed cellular metabolism to support uncontrolled cell proliferation^1^. Cancer cells exhibit increased nutrient uptake from the tumor microenvironment and altered intracellular pathways to meet the elevated demand for ATP production and biosynthesis of macromolecules required for rapid cell growth^2^. The Warburg effect demonstrates the higher demand of cancer cells for glucose than normal cells, which forms the basis for FDG-PET/CT imaging of cancer. Because cancer cell metabolism is heavily influenced by systemic and microenvironmental nutrient availability, dietary and pharmacologic interventions that alter cancer metabolism can be used as an adjuvant to standard antineoplastic therapies to improve clinical outcomes^3,4^. The precise mechanisms, however, through which the metabolic vulnerabilities of cancer cells can be systematically exploited remain poorly understood, limiting the translation of these strategies into clinical practice. Addressing this gap is critical for developing innovative approaches to improve patient care and refine individualized treatment regimens.

Emerging evidence indicates that nutrient availability in the blood or tumor microenvironment of individual cancer patients significantly influences tumor progression, drug responses, and clinical outcomes^5,6^. In addition, dietary interventions induce differential cytotoxic responses to anticancer drugs in cancer versus normal cells^7^. To advance the concept of oncometabolic precision medicine, we developed CM-SLP (Cancer Metabolism-based Synthetic Lethality Platform), a high-throughput screening platform designed to explore metabolic vulnerabilities associated with altered nutrient conditions. CM-SLP is the first comprehensive resource to evaluate the synthetic lethality of hyper- or hypo-nutrient states with therapeutic compounds, providing clinically feasible, safe, and effective strategies. By screening a panel of 1,813 FDA-approved oncology and non-oncology drugs, CM-SLP identifies unexpected cytotoxic effects arising from altered concentrations of glucose, glutamine, and fatty acids. Our findings reveal that nutrient states differentially sensitize cancer cells, while sparing normal cells, to various drugs prescribed for hypertension, arrhythmia, diabetes, inflammation, cancer, and infectious diseases. Through detailed mechanistic studies, CM-SLP enables the systematic analysis of an individual cancer patient’s metabolic profile, offering tailored modifications to treatment regimens to improve clinical outcomes. Furthermore, CM-SLP elucidates the effects of nutrient states on signaling transduction and drug responses, highlighting synthetic lethal interactions between therapeutic compounds and specific metabolites. Furthermore, CM-SLP has the potential to mitigate cancer treatment-induced metabolic syndromes, such as diabetes, obesity, and hypertension, by identifying metabolite-dependent cytotoxic side effects associated with various non-oncology drugs. Collectively, CM-SLP establishes a framework for personalized treatment regimens that reduce adverse effects while enhancing therapeutic efficacy in cancer patients with diverse metabolic states.

In this study, we further validate key findings from CM-SLP using both in vitro and in vivo models. Specifically, we demonstrate that candidate drugs from the CM-SLP^glu^ panel, including the antiarrhythmic drug propafenone and the antidiabetic drug biguanides, exhibit significant cytotoxicity and signaling alterations under hypoglycemic conditions. Mechanistic studies reveal that the therapeutic efficacy of these drug-nutrient combinations stems from targeted disruption of specific signaling pathways: propafenone/hypoglycemia suppresses tumor growth via inhibition of the LKB1-AMPK-TSC2-mTOR-S6K axis, while biguanides/hypoglycemia inhibit energy production through the FAK-Rho GTPase-LATS1/2-YAP/TAZ-TEAD axis. Therefore, we provide the rationale for substituting these metabolic conditions with targeted therapies, either mTOR or TEAD inhibitors. We further validated the synergistic effects of fenbendazole, niflumic acid, and tolfenamic acid with either hyperglycemic or hypo-glutamine conditions.

Together, CM-SLP presents a comprehensive framework for uncovering metabolite-dependent drug repurposing strategies and emphasizing the critical importance of evaluating drug responses in the context of nutrient availability. Our findings advance the current concept, add a substantial layer of patient-relevant complexity to drug treatments, and offer exciting new inroads to personalized medicine based on the metabolic status of individual cancer patients.

## Results

### Metabolic vulnerability landscape for non-oncology drugs according to nutrient availability

A panel of bioactive compounds, mostly FDA-approved drugs, was analyzed using CM-SLP to identify synthetic lethality between commonly prescribed medications and specific metabolite concentrations. This provided a comprehensive Resource for selecting clinically relevant drug-metabolite interactions. Through a high-throughput screening system, 1,813 FDA-approved compounds prescribed in diverse medical fields, including infectious disease (18.64%), endocrinology (10.42%), oncology (10.26%), cardiology (9.6%), neurology (9.21%), psychiatry (5.41%), immunology (5.36%), and others, were evaluated for metabolite-dependent cytotoxicity in A375 melanoma cells and their normal counterparts, HaCaT keratinocytes (**Fig. 1a,b**). These experiments were performed in culture medium containing high or low concentrations of critical metabolites, such as glucose, glutamine, and fatty acids. For the CM-SLP^glu^ platform, we used media containing 25 mM and 1 mM glucose (**Fig. 1c**); for the CM-SLP^gln^ platform, we used 2 mM and 0 mM glutamine (**Fig. 1d**); and for the CM-SLP^FA^ platform, we used normal versus charcoal-filtered serum, which has lowered lipid content (**Fig. 1e**). The nutrient content of these media, used to evaluate the metabolite-dependent cytotoxic effects of various drugs, reflects hyper- and hypo-nutrient statuses in either plasma or tumor interstitial fluid (TIF)^8^. The range for each CM-SLP quadrant was determined based on the correlation coefficients (r) and differential cytotoxicity against cancer and normal cells. Notably, lower correlation coefficients in cancer cells indicate that their metabolic rewiring renders them more vulnerable to various drugs compared to normal cells. Drugs in the first (group 1, upper right) and third quadrants (group 3, lower left) demonstrate minimal and maximal cytotoxicity, respectively, regardless of metabolite concentration. In contrast, the second (group 2, upper left) and fourth quadrants (group 4, lower right) contained compounds exhibiting metabolite concentration-dependent synthetic lethality, highlighting the significant influence of nutrient availability on the cytotoxicity of these drugs (**Figure 1c-e**). A substantial proportion of drugs located in groups 2 and 4 showed aberrant cytotoxicity under altered metabolite concentrations, specifically glucose (group 2, 12.08%; group 4, 2.14%), glutamine (group 2, 3.59%; group 4, 4.19%), and fatty acid (group 2, 4.14%; group 4, 3.86%). Among these drugs, essential subsets elicited cancer cell-specific cytotoxicity under modified nutrient conditions, including glucose (group 2, 23.64%; group 4, 22.22%), glutamine (group 2, 60%; group 4, 82.89%), and fatty acid (group 2, 69.33%; group 4, 88.57%) (Fig. 1f-h). To our surprise, many compounds in groups 2 and 4 that elicited both metabolite-dependent and cancer cell-specific synthetic lethality were non-oncology drugs. These results underscore the potential of CM-SLP to identify metabolite-dependent repurposing opportunities for a broad spectrum of non-oncology drugs, offering novel avenues for therapeutic innovation.

**Fig. 1.**
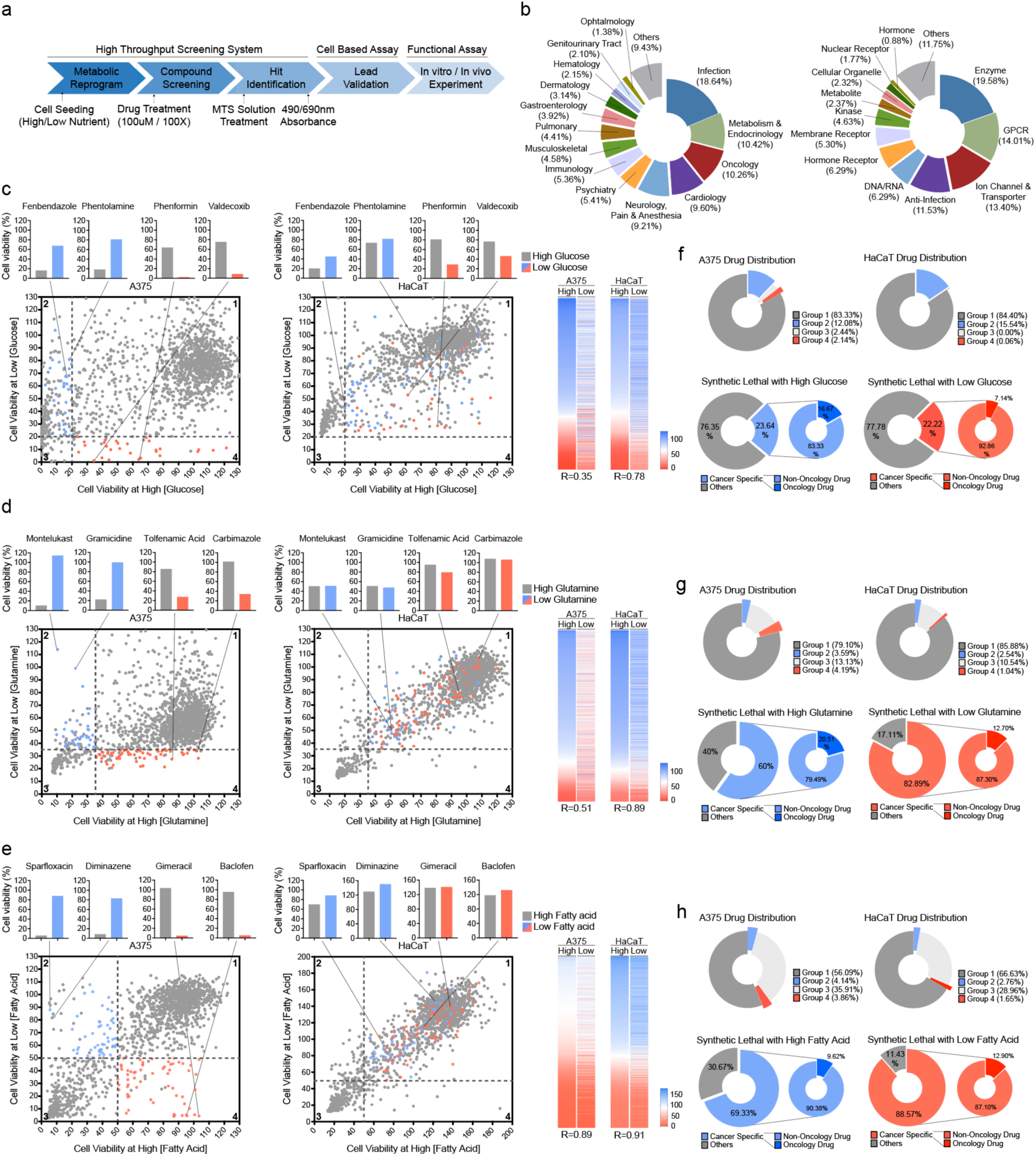
Landscape of the metabolic vulnerabilities of non-oncology drugs according to nutrient availability. **a** Schematic overview of the high-throughput CM-SLP screening system. 1,813 bioactive compounds were added to A375 melanoma and HaCaT keratinocyte cells under high or low nutrient concentrations, and cell viability was measured for each drug/nutrient status combination. **b** Composition of the CM-SLP drug library sorted by drug indications (left) and drug targets (right). **c-e** Dot plot summarizing the screen results from CM-SLP^glu^, CM-SLP^gln^, and CM-SLP^FA^ performed with the A375 (left panels) and HaCaT (right panels) cell lines. X-axis, relative cell viability at high nutrient concentrations; y-axis, relative cell viability at low nutrient concentrations. The dot plots are divided into 4 quadrants (group 1, upper right; group 2, upper left; group 3, lower left; and group 4, lower right) that distinguish drugs according to their cancer cell specificity. Blue dots, cancer cell-specific group 2 drugs; red dots, cancer cell-specific group 4 drugs. The bar graphs above the panels show cell viabilities of representative group 2 and group 4 candidate drugs. Y-axis, relative cell viability at high/low nutrient concentrations (left). Heatmap of CM-SLP and the correlation coefficient values (R) for each platform performed with A375 and HaCaT cells (right). **f-h** Analysis of CM-SLP candidates from CM-SLP^glu^, CM-SLP^gln^, and CM-SLP^FA^ performed with A375 and HaCaT cells. Drug composition of the CM-SLP results by quadrant (top row). The percentages of cancer-specific cytotoxic drugs in groups 2 and 4, as well as the percentage of non-oncology versus oncology drugs among the cancer-specific drugs, are shown for each CM-SLP screen performed with A375 cells (bottom row).

Representative compounds are shown above each CM-SLP panel (**Fig. 1c-e**). CM-SLP^glu^ revealed 48 compounds with cancer cell-specific synthetic lethality in the presence of high glucose (blue dots, group 2) and 28 compounds for low glucose (red dots, group 4). Notable examples include the anti-parasitic drug fenbendazole and the non-selective α-adrenergic antagonist phentolamine, which showed synthetic lethality in high glucose environments (group 2). Conversely, the anti-diabetic drug phenformin and the anti-inflammatory drug valdecoxib exhibited enhanced anticancer activity under low glucose conditions (group 4) (**Fig. 1c**). In the CM-SLP^gln^ platform, 39 compounds demonstrated cancer cell-specific synthetic lethality in high glutamine (blue dots, group 2) and 63 drugs in low glutamine (red dots, group 4) conditions. The asthma drug montelukast and the antimicrobial drug gramicidin exhibited synthetic lethality in the presence of high glutamine (group 2), whereas the anti-inflammatory drug tolfenamic acid and the anti-thyroid drug carbimazole displayed improved anticancer efficacy in low glutamine conditions (group 4) (**Fig. 1d**). In the CM-SLP^FA^ platform, 52 compounds were found to exhibit cancer cell-specific synthetic lethality in high fatty acid concentrations (blue dots, group 2), while 62 in low concentrations of fatty acids (red dots, group 4). Examples include the anti-bacterial drugs sparfloxacin and diminazene, which exhibited synthetic lethality in high concentrations of fatty acids (group 2), whereas the antineoplastic adjuvant gimeracil and the muscle spasticity drug baclofen (group 4) showed anticancer effects in low concentrations of fatty acids (**Fig 1e**). Collectively, these findings underscore how nutrient availability significantly influences therapeutic responses of non-oncology drug. The CM-SLP candidates provides valuable insights into designing novel synthetic lethal combination therapies that target cancer metabolism, offering a foundation for innovative and metabolically informed therapeutic strategies.

### Nutrient availability-dependent anticancer drug repurposing opportunities

The CM-SLP score represents the ratio between cell viability in the context of high versus low metabolite concentrations. CM-SLP scores were calculated by the following equation: CM-SLP score = log0.5([cell viability at high nutrient]/[cell viability at low nutrient]) (**Supplementary Tables 1-3**). Compounds with significantly high or low CM-SLP scores were identified from each metabolite screening platform as candidates whose cancer cell-specific cytotoxic effects were sensitive to nutrient availability. Among the top candidates identified in each CM-SLP screen, CM-SLP^glu^ had the lowest R-value and therefore the most compounds with significant CM-SLP scores (**Fig. 2a, b**). Although hypo-nutrient conditions commonly enhance cytotoxicity, CM-SLP^glu^ group 2 drugs exhibited synergistic effects under hyperglycemic conditions^1^. These group 2 compounds, primarily prescribed for infectious diseases, endocrinology, and oncology target DNA/RNA, ion channels/transporters, and infections (**Fig. 2c**). 23% of CM-SLP^glu^ group 2 drugs interfered with DNA replication by targeting DNA intercalation and topoisomerases, suggesting that the accelerated cell cycle progression characteristic of cancer cells under hyperglycemic conditions may confer increased vulnerability to these agents. Importantly, hyperglycemia is a common side effect of anticancer treatments and a well-established risk factor for cancer development and relapse among survivors^9,10^. Therefore, CM-SLP^glu^ group 2 candidate drugs may hold particular therapeutic value for cancer patients experiencing metabolic syndromes, such as diabetes, insulin resistance, or obesity, offering a targeted approach to address both metabolic and oncologic challenges.

**Fig. 2.**
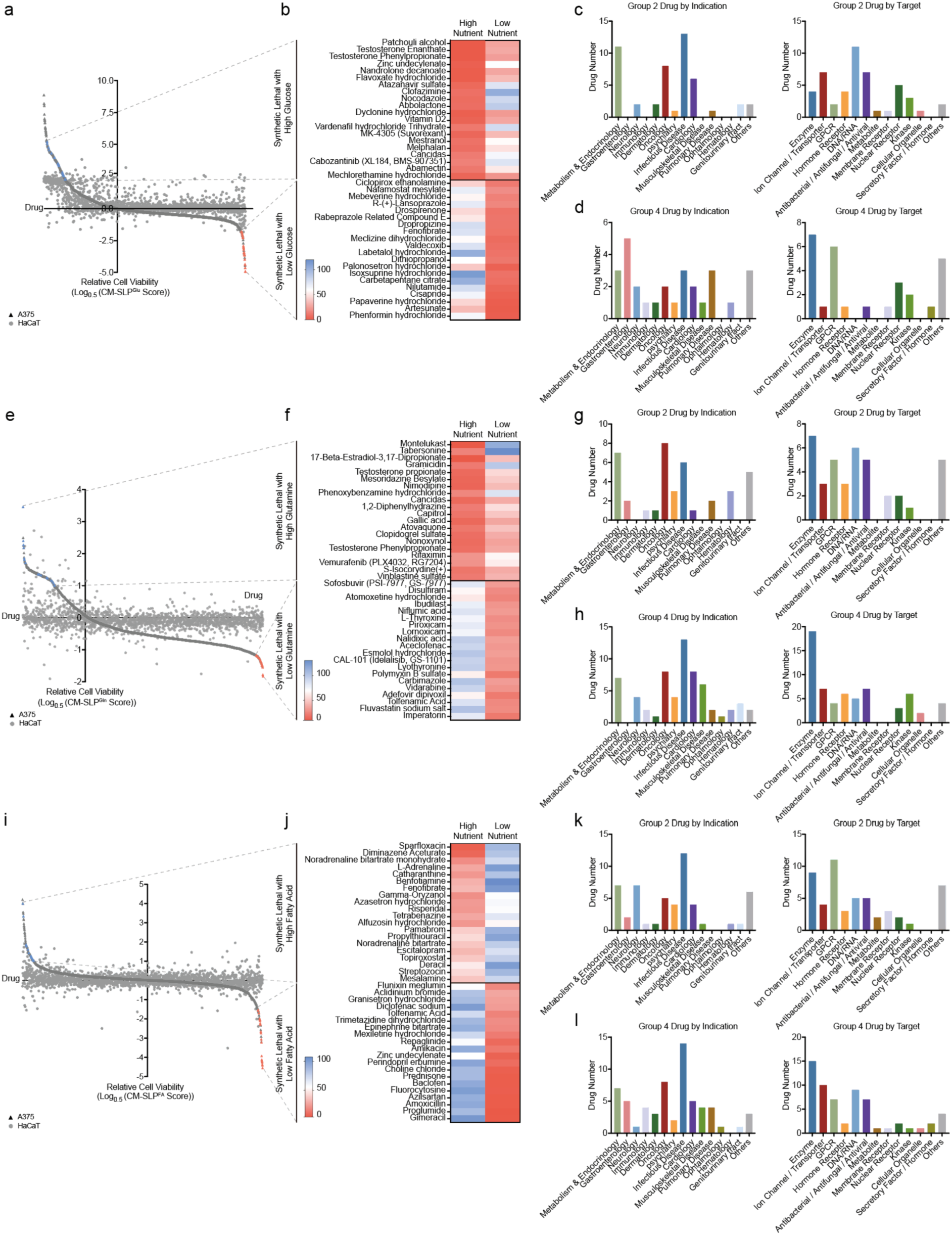
Nutrient availability-dependent anticancer drug repurposing opportunities. **a** High-throughput screening results for the CM-SLP^glu^ panels. Drugs are listed according to their individual top CM-SLP scores (left). X-axis, dot plot of individual compounds; y-axis, CM-SLP scores; blue dots, cancer cell-specific group 2 drugs; red dots, cancer cell-specific group 4 drugs; black dots, CM-SLP results performed with A375 cells; gray dots, CM-SLP results performed with HaCaT cells. **b** Heatmap of top CM-SLP candidates that show significant CM-SLP scores from groups 2 and 4 of the CM-SLP^glu^ screens. **c** Indications and molecular target analyses of group 2 CM-SLP candidates from the CM-SLP^glu^ screens. **d** Indications and molecular target analyses of group 4 CM-SLP candidates from the CM-SLP^glu^ screens. **e** High-throughput screening results for the CM-SLP^gln^ panels. Drugs are listed according to their individual top CM-SLP scores (left). X-axis, dot plot of individual compounds; y-axis, CM-SLP scores; blue dots, cancer cell-specific group 2 drugs; red dots, cancer cell-specific group 4 drugs; black dots, CM-SLP results performed with A375 cells; gray dots, CM-SLP results performed with HaCaT cells. **f** Heatmap of top CM-SLP candidates that show significant CM-SLP scores from groups 2 and 4 of the CM-SLP^gln^ (**f**) screens. **g** Indications and molecular target analyses of group 2 CM-SLP candidates from the CM-SLP^gln^ screens. **h** Indications and molecular target analyses of group 4 CM-SLP candidates from the CM-SLP^gln^ screens. **i** High-throughput screening results for the CM-SLP^FA^ panels. Drugs are listed according to their individual top CM-SLP scores (left). X-axis, dot plot of individual compounds; y-axis, CM-SLP scores; blue dots, cancer cell-specific group 2 drugs; red dots, cancer cell-specific group 4 drugs; black dots, CM-SLP results performed with A375 cells; gray dots, CM-SLP results performed with HaCaT cells. **j** Heatmap of top CM-SLP candidates that show significant CM-SLP scores from groups 2 and 4 of the CM-SLP^FA^ screens. **k** Indications and molecular target analyses of group 2 CM-SLP candidates from the CM-SLP^FA^ screens. **l** Indications and molecular target analyses of group 4 CM-SLP candidates from the CM-SLP^FA^ screens.

In CM-SLP^glu^, representative group 4 drugs with significant CM-SLP scores and cancer specificity included FDA-approved drugs prescribed in the fields of gastroenterology, endocrinology and pulmonary diseases. These drugs target a range of biomolecules, such as enzymes, ion channels/transporters, and nuclear receptors (**Fig. 2d**). Synergism with hypoglycemia could be achieved through dietary interventions like caloric restriction (CR) or intermittent fasting (IF), both of which enhance the efficacy of anticancer treatments due to differential responses of cancer and normal cells to metabolic stress^4,7^. In addition, glucose-lowering agents, such as GLUT inhibitors, SGLT2 inhibitors, or GLP-1 receptor agonists, could exhibit synthetic lethality when paired with group 4 drugs. Mechanistically, several group 4 drugs, such as phenformin, nilutamide, and valdecoxib are known to target mitochondrial functions. This suggests blockade of the metabolic shift between glycolysis and oxidative phosphorylation (OXPHOS) may account for the increased vulnerability of cancer cells to these combination treatments^11–13^.

In CM-SLP^gln^, group 2 compounds with significant CM-SLP scores and cancer specificity are mostly prescribed in the fields of oncology, endocrinology, and infectious diseases. These drugs target a diverse range of molecular pathways and entities, including enzymes, G-protein-coupled receptors (GPCRs), DNA/RNA, and various microroganisms (**Fig. 2e-g**). The CM-SLP^gln^ group 4 compounds with significant CM-SLP scores included drugs prescribed in the fields of oncology, cardiology, and infectious diseases. These compounds are notably enriched for drugs that specifically target enzymes, suggesting a pivotal role for enzymatic pathways in their mechanism of action under altered glutamine conditions. These findings highlight the broad therapeutic potential of metabolite-dependent drug repurposing across multiple clinical disciplines (**Fig. 2h**).

In CM-SLP^FA^, the group 2 compounds with high CM-SLP scores and cancer specificity included drugs mostly prescribed for the treatment of infectious diseases, metabolic or endocrine disorders, and neurological conditions. These drugs are enriched for drugs targeting GPCRs and enzymes (**Fig. 2i-k**). Interestingly, many of the group 2 CM-SLP^FA^ drugs induce DNA damage or DNA fragmentation, suggesting that these compounds selectively increase the vulnerability of cancer cells with higher rates of DNA replication than normal cells ^14–16^. As with hyperglycemia, high fatty acid concentrations lead to the over-production of proliferation signals. Hyperlipidemia, a condition often linked to increased cancer risk^9,17^, further underscores the relevance of group 2 candidates for simultaneously targeting cancer progression and managing comorbid metabolic disorders. The group 4 CM-SLP^FA^ compounds with significant CM-SLP scores and cancer selectivity included drugs typically prescribed for the treatment of infectious diseases, oncology, and metabolic and endocrine disorders. They included drugs targeting enzymes, ion channels, and transporters, demonstrating their broad mechanistic potential in addressing cancer-specific vulnerabilities under low fatty acid conditions (**Fig. 2l**).

Together, we identified the top CM-SLP candidates exhibiting high cytotoxicity under altered metabolite concentrations, offering new opportunities for repurposing non-oncology drugs by incorporating the metabolic status of individual patients. Although further clinical validation will be necessary, we propose that group 2 CM-SLP candidates hold promise for treating cancer patients with elevated blood nutrient levels or metabolic syndromes. Conversely, group 4 CM-SLP candidates may be particularly effective when combined with dietary interventions or blood metabolite-lowering agents, providing a tailored approach to enhance therapeutic outcomes.

### Nutrient-sensitive cytotoxicity driven via cancer-related signaling perturbations

Since the CM-SLP^glu^ screen showed the lowest R-value, identifying glucose as the critical metabolite responsible for metabolite-sensitive drug responses, we proceeded to validate the drugs classified under CM-SLP^glu^ drugs from groups 1 through 4. This validation was based on two criteria: their CM-SLP score and specificity towards cancer cells. We altered glucose levels in the culture media to contrast high (25 mM) and low (1 mM) glucose concentrations, and we used a GLUT inhibitor to impair glucose uptake. As expected, a group 1 drug, warfarin, did not elicit cytotoxicity under either high or low glucose concentrations (**Fig. 3a**). In contrast, the group 2 drug fenbendazole elicited significant cytotoxicity in the presence of high glucose, but not in glucose-restricted conditions (**Fig. 3b**). To our knowledge, this is the first demonstration of hyperglycemia conferring cytotoxicity on a non-oncology drug. On the other hand, group 4 drugs, including phenformin, valdecoxib, verapamil, and nilutamide, elicited minimal cytotoxicity in high glucose conditions but synergistic cytotoxicity in low glucose or in the presence of a GLUT inhibitor (**Fig. 3c-f**). These CM-SLP candidates confirm the notion that nutrient availability can be a critical determinant of cytotoxic efficacy against cancer cells. Next, to determine the physiological relevance of the *in vitro* glucose sensitivity, we tested various glucose concentrations that are feasible to achieve in human patients. Normal glucose level was set to 5 mM, hypoglycemia to < 5 mM, and hyperglycemia to > 5 mM. Remarkably, glucose levels below 5 mM were enough to induce propafenone and phenformin cytotoxicity, whereas glucose levels above 5 mM were sufficient to induce fenbendazole cytotoxicity (**Fig. 3g-h**). These results demonstrate that the range of high and low glucose concentrations that confer cytotoxicity to these non-oncology drugs *in vitro* are likely to be physiologically relevant *in vivo*.

**Fig. 3.**
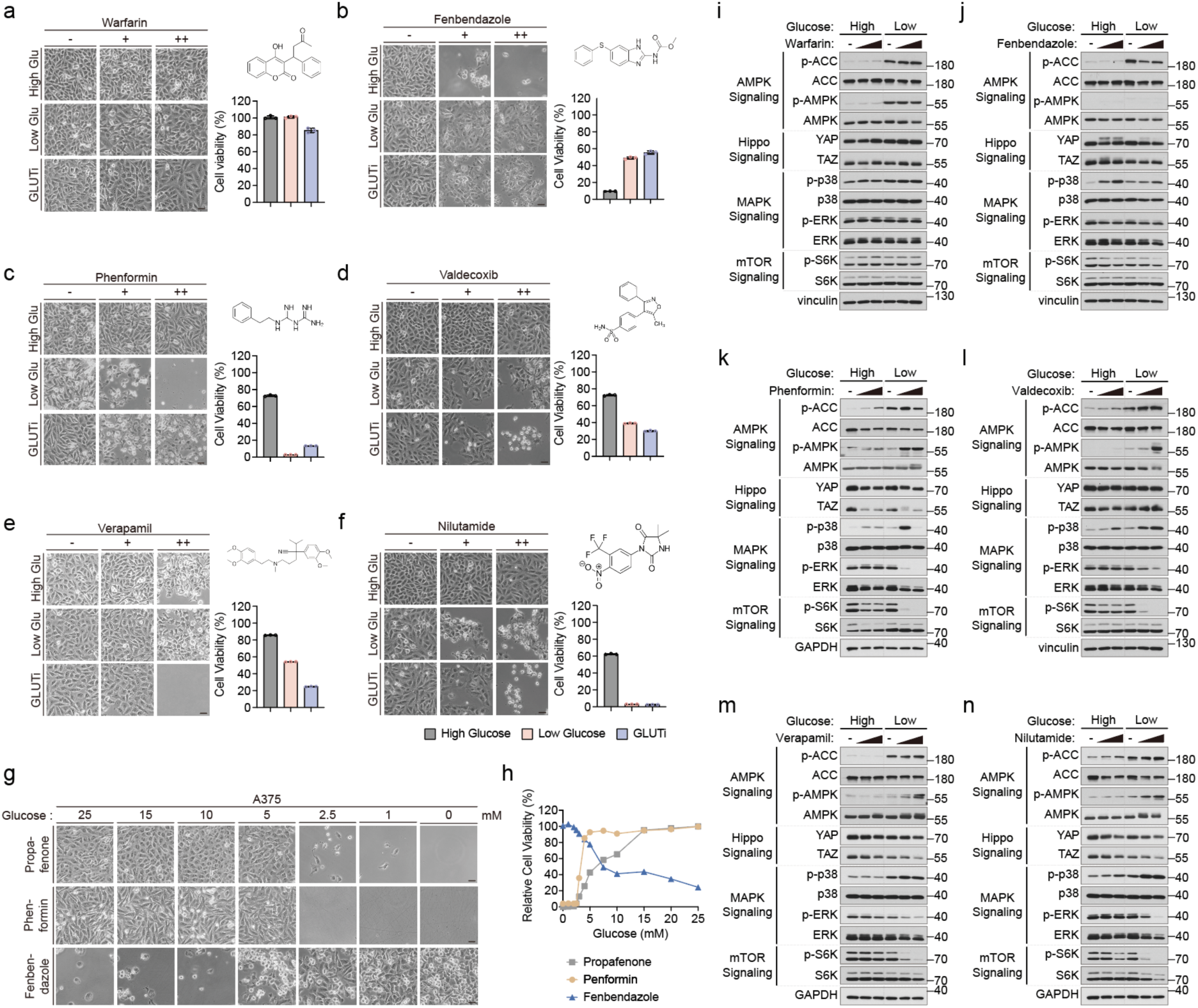
CM-SLP candidates induce glucose concentration-dependent cytotoxicity and signaling perturbations. **a** Viability of A375 cells subjected to warfarin (group 1; 0, 50, 100 μM, 8 h) treatment in the presence of high glucose (25 mM), low glucose (1 mM, 11 h), or the GLUT inhibitor BAY-876 (10 mM). **b** Viability of A375 cells subjected to fenbendazole (group 2; 0, 50, 100 μM, 8 h) treatment in the presence of high glucose (25 mM), low glucose (1 mM, 11 h), or the GLUT inhibitor BAY-876 (10 mM). **c-f** Viability of A375 cells subjected to treatment with group 4 drugs in the presence of high glucose (25 mM), low glucose (1 mM, 11 h), or the GLUT inhibitor BAY-876 (10 mM). Each CM-SLP drug was administered at 50 μM or 100 μM (8 h) to the A375 cell line. Drug structure (right top). Relative cell viability for each drug under high/low glucose concentrations (right bottom). **g, h** Viability of A375 cells treated with various glucose concentrations (0, 1, 2.5, 5, 10, 15, and 25 mM, 11 h) combined with either propafenone (30 μM, 24 h), phenformin (100 μM, 8 h), or fenbendazole (12.5 μM, 8 h). Anticancer efficacy of propafenone and phenformin was apparent in < 5 mM glucose, while it appeared for fenbendazole in > 5 mM glucose. **i-n** Immunoblotting analysis of cancer-related signaling pathways in A375 cells treated with warfarin (**i**), fenbendazole (**j**), phenformin (**k**), valdecoxib (**l**), verapamil (**m**), and nilutamide (**n**) combined with either high (25 mM) or low (1 mM, 11 h) glucose. -, negative control; low dose, 50 μM; high dose, 100 μM. All drug treatments lasted 8 h.

Because it is likely that cancer-related signaling pathway perturbations induced by altered metabolite concentrations contribute to the synthetic lethality of CM-SLP candidates, we next looked for changes in AMPK, Hippo, MAPK, and mTOR signaling pathways in melanoma cells exposed to different glucose concentrations. Consistent with our cell viability results, the group 1 CM-SLP^glu^ drug, warfarin, failed to further alter any of the signaling pathways we examined under both high and low glucose concentrations (**Fig. 3i**). We found, however, that the group 2 drug fenbendazole altered YAP/TAZ and p38 under high glucose conditions (**Fig. 3j**), and that the group 4 drugs phenformin, valdecoxib, verapamil, and nilutamide altered AMPK, mTOR, p38, YAP/TAZ, and ERK signaling under low glucose conditions (**Fig. 3k-n**). Interestingly, these compounds are commonly known to induce mitochondrial dysfunction. This may explain their synergism with hypoglycemic conditions by disrupting energy sensing and growth signaling pathways.

To further investigate the potential of drug re-purposing based on metabolite availability, we analyzed the anti-cancer efficacy of FDA-approved drugs, both those originally approved for oncology purposes and those approved for non-oncology indications. Among the CM-SLP^glu^ candidates, we selected fenbendazole (group 2) and phenformin (group 4) as representative non-oncology drugs affected by high or low glucose concentrations and measured their anticancer efficacy against oncology drugs prescribed to melanoma and NSCLC patients. Oncology drugs within the FDA-approved drug panel were distributed across groups 1 through 4 (**Supplementary Fig. 1a, c**). The MAPK pathway inhibitors dabrafenib (BRAF inhibitor) and trametinib (MEK1 inhibitor) appeared among the group 1 CM-SLP^glu^ drugs, which were less dependent on glucose concentrations (**Supplementary Fig. 1b**). The group 2 drug fenbendazole and the group 4 drug phenformin, however, evoked dramatic dose-dependent cancer cell death in low and high glucose conditions, respectively (**Supplementary Fig. 1d**). Moreover, it was apparent that, under similar time courses and drug concentrations, phenformin/low glucose and fenbendazole/high glucose conditions exerted enhanced cytotoxicity compared to the MAPK inhibitors. We further confirmed the glutamine sensitivity of the CM-SLP^gln^ group 4 non-oncology drugs niflumic acid and tolfenamic acid that could be re-positioned as anticancer agents depending on glutamine availability of cancer cells (**Supplementary Fig. 2a-c**). These results suggest that nutrient availability can be a critical factor influencing the cytotoxicity of commonly prescribed non-oncology drugs by dysregulating cancer-related signaling pathways.

### Propafenone/low glucose treatment promotes cancer cell death via the AMPK-mTOR-S6K axis

Cardiovascular disease (CVD) remains one of the leading causes of mortality worldwide. The intersection between CVD and anti-cancer treatments is an emerging concern, because anti-cancer treatment often have potential side effects that can induce cardiovascular complications^18^. In such cases, anticancer treatments can inadvertently cause cardiac rhythm disorders, such as arrhythmia^19,20^. Numerous studies indicate that CVD patients may also face a heightened risk of developing cancer, emphasizing the need for an integrated therapeutic approach that considers both CVD and cancer in a unified framework^21,22^. Because both CVD and cancer are associated with significant metabolic alterations^23^, we aimed to investigate the metabolic association between cancer and CVD. Here we focused on propafenone, an anti-arrhythmic drug in CM-SLP^glu^ group 4, as a representative non-oncology drug with significantly enhanced cytotoxicity in hypoglycemic conditions.

To explore the anti-cancer efficacy of propafenone under glucose-restricted conditions, we tested the cytotoxic effect of propafenone/hypoglycemia treatment in a wide range of cancer types. In addition to A375 melanoma cells, we tested 786-O renal cancer cells, H358 lung cancer cells, and BT549 breast cancer cells, along with their normal counterparts, HEMn primary epidermal melanocytes, HREC primary renal epithelial cells, BEAS-2B non-tumorigenic lung epithelial cells, and MCF-10A non-malignant breast epithelial cells. Under glucose-restricted conditions, propafenone exerted dramatic cytotoxicity in all cancer cell lines (**Fig. 4a-d**) but not in their corresponding non-cancerous counterparts (**Fig. 4e-h**). This validates the anti-cancer effect of combined propafenone/hypoglycemia treatment.

**Fig. 4.**
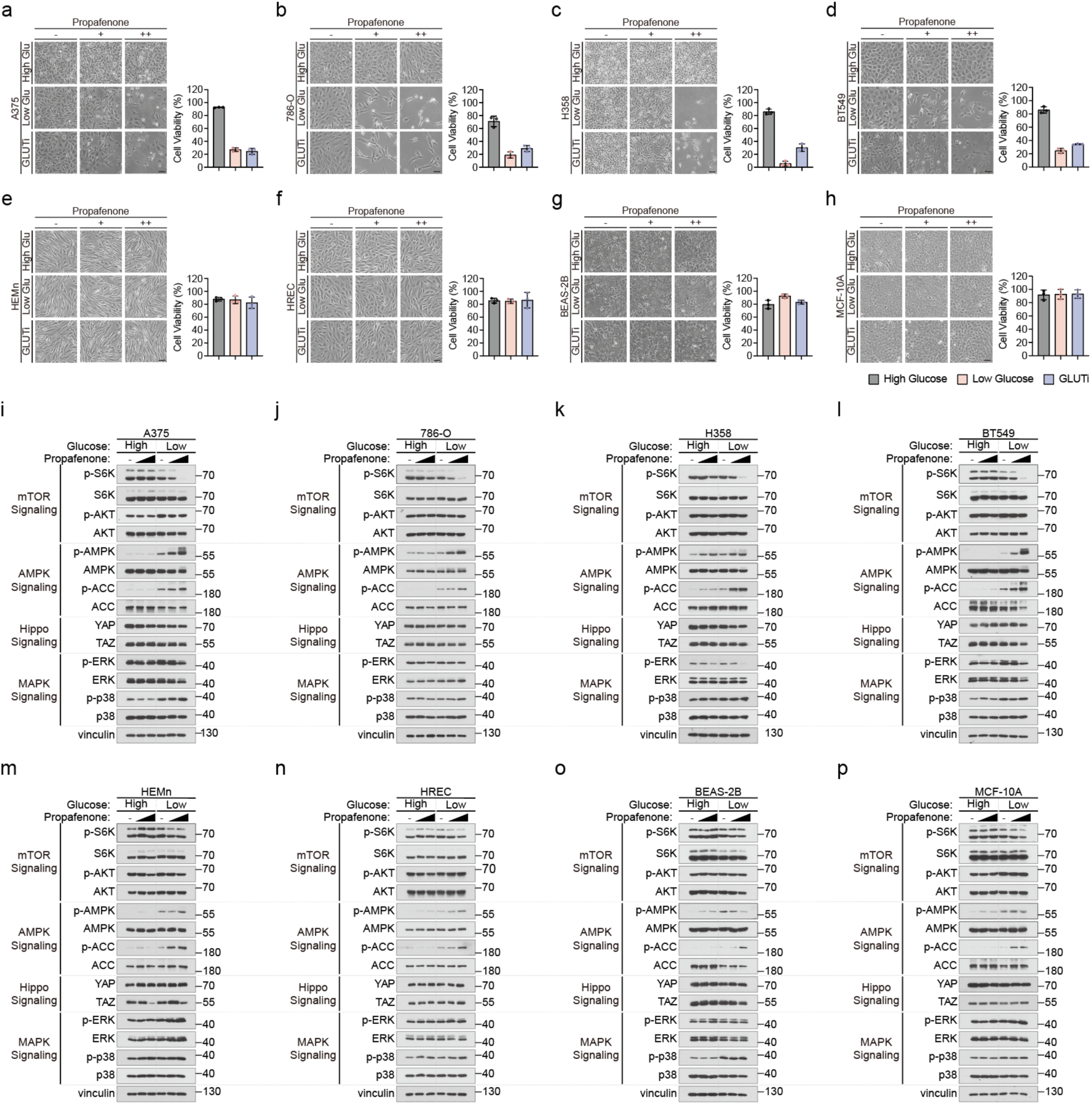
Propafenone/low glucose treatment alters oncogenic signaling pathways in cancer cells. **a** Viability of A375 cells subjected to propafenone (0, 10, 30 μM, 24 h) treatment in the presence of high glucose (25 mM), low glucose (1 mM), or the GLUT inhibitor BAY-876 (10 μM). Relative viability for each cell type under high/low glucose concentrations (right bottom). **b** Viability of 786-O cells subjected to propafenone (0, 10, 30 μM, 48 h) treatment in the presence of high glucose (25 mM), low glucose (1 mM), or the GLUT inhibitor BAY-876 (3 μM). Relative viability for each cell type under high/low glucose concentrations (right bottom). **c** Viability of H358 cells subjected to propafenone (0, 10, 30 μM, 24 h) treatment in the presence of high glucose (25 mM), low glucose (1 mM), or the GLUT inhibitor BAY-876 (10 μM). Relative viability for each cell type under high/low glucose concentrations (right bottom). **d** Viability of BT549 cells subjected to propafenone (0, 10, 30 μM, 24 h) treatment in the presence of high glucose (25 mM), low glucose (1 mM), or the GLUT inhibitor BAY-876 (7 μM). Relative viability for each cell type under high/low glucose concentrations (right bottom). **e** Viability of HEMn cells subjected to propafenone (0, 10, 30 μM, 24 h) treatment in the presence of high glucose (25 mM), low glucose (1 mM), or the GLUT inhibitor BAY-876 (10 μM). Relative viability for each cell type under high/low glucose concentrations (right bottom). **f** Viability of HREC cells subjected to propafenone (0, 10, 30 μM, 48 h) treatment in the presence of high glucose (25 mM), low glucose (1 mM), or the GLUT inhibitor BAY-876 (3 μM). Relative viability for each cell type under high/low glucose concentrations (right bottom). **g** Viability of BEAS-2B cells subjected to propafenone (0, 10, 30 μM, 24 h) treatment in the presence of high glucose (25 mM), low glucose (1 mM), or the GLUT inhibitor BAY-876 (10 μM). Relative viability for each cell type under high/low glucose concentrations (right bottom). **h** Viability of MCF-10A cells subjected to propafenone (0, 10, 30 μM, 24 h) treatment in the presence of high glucose (25 mM), low glucose (1 mM), or the GLUT inhibitor BAY-876 (7 μM). Relative viability for each cell type under high/low glucose concentrations (right bottom). **i-p** Immunoblotting analysis of cancer-related signaling pathways in cells treated with propafenone combined with either high (25 mM) or low (1 mM) glucose. -, negative control; low dose, 10 μM; high dose, 30 μM. A375, HEMn, H358, BES-2B, BT549 and MCF-10A were treated for 16 h. 786-O, HREC cells were treated for 8 h.

To elucidate the signaling pathways underlying the glucose-dependent differential cytotoxicity of propafenone, we conducted a comprehensive analysis of cancer-related signaling mechanisms. Our results revealed distinct patterns of S6K and AMPK phosphorylation between cancer cells and their counterparts. Notably, the combination of propafenone and low glucose induced significant dephosphorylation of S6K in cancer cells, indicating robust inhibition of the mTORC1 activity under these conditions^24^. In contrast, AKT phosphorylation, a marker of mTORC2 activity, remained unchanged, suggesting mTORC2 signaling was unaffected. Concurrently, we observed increased AMPK phosphorylation, which indicates energy stress within the cellular environment (**Fig. 4i-l**)^25^. In the non-cancerous counterparts, we observed minimal S6K dephosphorylation and negligible AMPK phosphorylation, indicating that the energy stress response was elicited predominantly in the cancer cells (**Fig. 4m-p**). Collectively, these findings underscore the pivotal role of the AMPK-mTORC1 signaling axis in mediating the differential cytotoxicity between cancer and non-cancer cells. This differential response highlights the therapeutic potential of targeting this pathway to exploit the metabolic vulnerabilities of cancer cells while sparing normal cells.

### Redirecting metabolic interventions toward mTOR-targeted therapies improves the clinical applicability of CM-SLP

Given that TSC2 inhibits mTORC1 when AMPK is phosphorylated^26^, we performed experiments to determine how TSC2 influences the observed cytotoxic effects of propafenone and low glucose treatment. Using p53 knockout (KO) MEF cells, p53/TSC2 double KO MEF cells, and TSC2-deficient hepatocellular carcinoma cell lines (SNU886 and SNU878), we observed that TSC2 status plays a pivotal role in determining treatment efficacy. Specifically, TSC2 wild-type MEF cells exhibited a robust cytotoxic response under propafenone and low glucose treatment, as evidenced by significant AMPK phosphorylation and S6K dephosphorylation (**Fig. 5a, b**). In contrast, TSC2 KO MEF cells, as well as the TSC2-deficient SNU886 and SNU878 cell lines, were insensitive to combined treatment (**Fig. 5c-e**). TSC mutant cells showed AMPK phosphorylation but failed to exhibit S6K dephosphorylation (**Fig. 5f-h**). These findings suggest that the absence of TSC2 disrupts the downstream inhibition of mTORC1, underscoring the necessity of the TSC2-mTORC1 axis for mediating the cytotoxic effects of propafenone/hypoglycemia treatment.

**Fig. 5.**
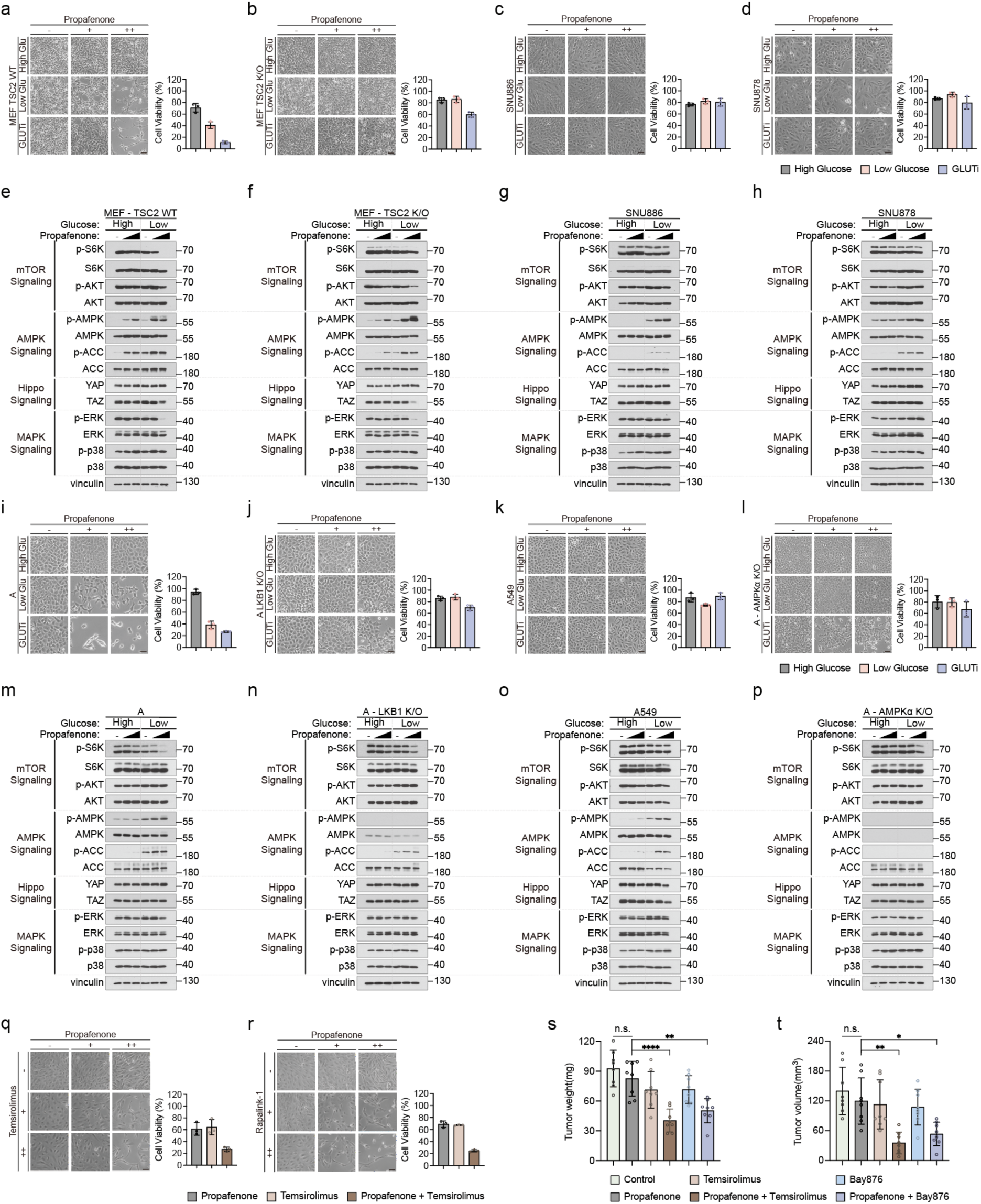
LKB1-AMPK-TSC2-mTOR pathway-mediated propafenone/low glucose-induced anticancer efficacy. **a-d** Viability of all cells subjected to propafenone (0, 5, 15 μM, 24 h) treatment in the presence of high glucose (25 mM), low glucose (1 mM), or the GLUT inhibitor BAY-876 (2 μM). Relative viability for each cell type under high/low glucose concentrations (right bottom). **e-h** Viability of A cells subjected to propafenone (0, 10, 20 μM, 24 h) treatment in the presence of high glucose (25 mM), low glucose (1 mM), or the GLUT inhibitor BAY-876 (10 μM). Viability of A549 cells subjected to propafenone (0, 10, 30 μM, 24 h) treatment in the presence of high glucose (25 mM), low glucose (1 mM), or the GLUT inhibitor BAY-876 (10 μM). Relative viability for each cell type under high/low glucose concentrations (right bottom). **i-l** Immunoblotting analysis of cancer-related signaling pathways in cells treated with propafenone combined with either high (25 mM) or low (1 mM) glucose. -, negative control; low dose, 5 μM; high dose, 15 μM. All cells were treated for 4 h. **m-p** Immunoblotting analysis of cancer-related signaling pathways in cells treated with propafenone combined with either high (25 mM) or low (1 mM) glucose. -, negative control; low dose, 10 μM; high dose, 20 μM. Each group of HEK293A cells was treated for 2 h, while A549 cells were treated for 16 h. **q** Selective cytotoxicity of propafenone with the mTOR inhibitor temsirolimus. Propafenone was administered at 10 μM or 30 μM, while temsirolimus was administered at 100 nM or 300 nM. Both drugs were administered for 72 h at low cell density. Relative viability for each cell type treated with propafenone, temsirolimus, or the combination of propafenone and temsirolimus (right bottom). **r** Selective cytotoxicity of propafenone with the mTOR inhibitor rapalink-1. Propafenone was administered at 10 μM and 30 μM, while rapalink-1 was administered at 1 nM and 3 nM. Both drugs were administered for 72 h at low cell density. Relative viability for each cell type treated with propafenone, rapalink-1, or the combination of propafenone and rapalink-1 (right bottom). **s, t** *In vivo* xenograft tumor growth measured by tumor weight (mg) and volume (mm^3^) after administration of the combinations of propafenone/temsirolimus or propafenone/BAY-876.

To further dissect the signaling pathways involved, we next turned our attention upstream to LKB1, which is a key regulator of AMPK^27^. We tested the cytotoxic effects of propafenone and low glucose treatment on wild-type HEK293A cells, LKB1-KO HEK293A cells, LKB1-deficient A549 lung carcinoma cells, and AMPK-KO HEK293A cells. While wild-type cells displayed significant cell death accompanied by AMPK phosphorylation and S6K dephosphorylation under treatment (**Fig. 5i, j**), cells lacking LKB1, such as LKB1-KO cells, A549 cells, and AMPK-KO HEK293A cells, showed markedly reduced cytotoxic responses (**Fig. 5k-p**). Notably, these cells demonstrated insufficient AMPK phosphorylation and minimal S6K dephosphorylation compared to their wild-type counterparts. This result highlights the critical role of the LKB1-AMPK axis, which is required for the downstream inhibition of mTORC1 and subsequent cytotoxic effects.

Recognizing that dietary interventions such as hypoglycemia can pose challenges for patients with metabolic imbalances, we explored alternative therapeutic approaches using mTOR inhibitors to bypass the reliance on low glucose conditions. We tested temsirolimus, an FDA-approved mTOR inhibitor, and Rapalink-1, a third-generation mTOR inhibitor^28,29^, in combination with propafenone. Both combinations exhibited strong synergistic effects, significantly enhancing the anticancer efficacy of propafenone without the need for hypoglycemia (**Fig. 5q, r**). These findings suggest that integrating mTOR inhibitors into treatment regimens could overcome the challenges associated with dietary interventions, making the therapies more clinically applicable.

Several mTOR inhibitors, such as temsirolimus and everolimus, have been approved by the FDA for renal cell carcinoma (RCC) patients^30,31^. To validate these findings *in vivo*, we conducted experiments using NSG mice subcutaneously injected with 786-O renal cancer cells. After the tumors reached comparable size, the mice were randomly assigned to treatment groups and administered propafenone (30 mg/kg), temsirolimus (0.3 mg/kg), or the glucose uptake inhibitor BAY-876 (2 mg/kg)^32^, either alone or in combination, every other day for 30 days. Remarkably, the combinations of propafenone/BAY-876 and propafenone/temsirolimus significantly reduced tumor weight and volume compared to single treatments, with no significant changes in body weight (**Fig. 5s, t**). These results confirm the efficacy and safety of the combined treatments and demonstrate their potential to selectively target cancer cells without affecting overall health.

Collectively, these findings demonstrate that the anticancer effects of propafenone/low glucose treatment are mediated through the LKB1-AMPK-TSC2-mTORC1 axis. Furthermore, the observed synergy between propafenone and mTOR inhibitors offers a promising alternative strategy for cancer treatment, particularly for patients for whom dietary interventions are not feasible. By tailoring therapeutic approaches based on the signaling pathways elucidated through the CM-SLP framework and leveraging the unique metabolic vulnerabilities of cancer cells, these strategies pave the way for more effective and personalized oncology treatments.

### Propafenone targets the K_Ca_ channel to inhibit AMPK-mTOR axis and promote cancer cell death

To uncover the specific molecular targets associated with the therapeutic combination of propafenone and low glucose, we initially investigated the voltage-gated sodium (Na_v_) channel, which is the established target of propafenone in cardiac muscle cells. We found, however, that 786-O cells do not express Na_v_ channels (**Supplementary Fig. 3a**). Thus, we broadened our investigation to other potential antiarrhythmic drug targets. We examined the expression of voltage-gated calcium (Ca_v_) and potassium channels but were unable to detect either in 786-O cells (**Supplementary Fig. 3b**, **Fig. 6a**).

**Fig. 6.**
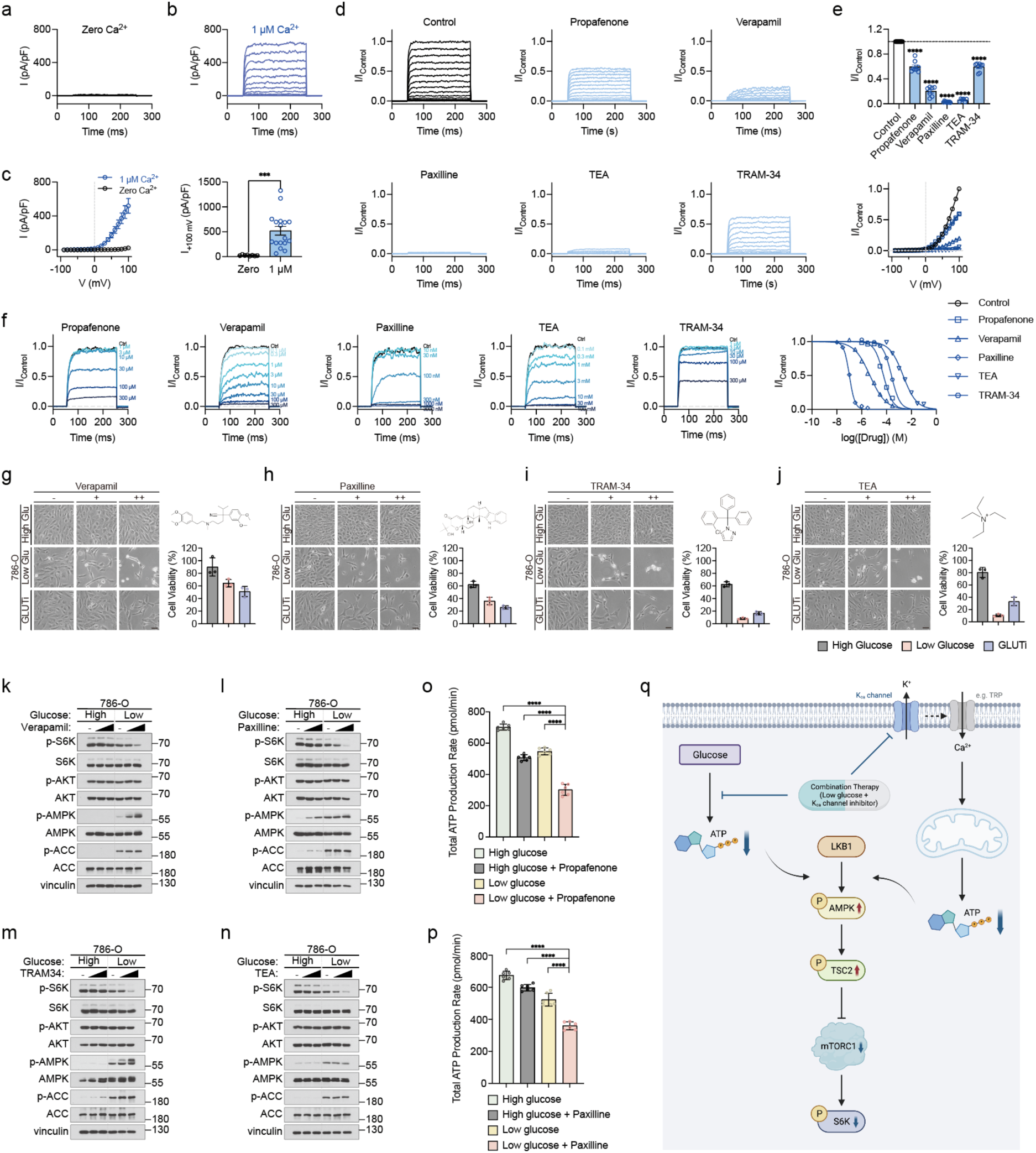
Propafenone targets the K_Ca_ channel to inhibit AMPK-mTOR axis and promote cancer cell death. **a** Representative traces of whole-cell K^+^ currents in 786-O cells in which intracellular Ca^2+^ was fixed at zero. **b** Representative traces of whole-cell K^+^ currents in 786-O cells in which intracellular Ca^2+^ was fixed at 1 μM. **c** Voltage-current relationship and current density of whole-cell K^+^ currents in zero or 1 μM intracellular Ca^2+^. Current density at 100 mV in 0 μM Ca^2+^, 24.33 ± 3.73 pA/pF (n = 8) or 1 μM Ca^2+^, 519.91 ± 87.22 pA/pF (n = 17). **d** Representative traces of normalized whole-cell K^+^ currents with or without chemicals. Intracellular Ca^2+^ was fixed at 1 μM, and chemicals were added to the extracellular side. Chemical concentrations: propafenone, 20 μM; verapamil, 20 μM; paxilline, 20 μM; TEA, 20 mM; and TRAM-34, 60 μM. **e** Normalized peak currents and voltage-current relationships for whole-cell K^+^ currents with or without chemicals. Normalized current at 100 mV with the following chemicals: propafenone, 0.594 ± 0.090 (n = 8); verapamil, 0.201 ± 0.029 (n = 7); paxilline, 0.029 ± 0.003 (n = 9); TEA, 0.060 ± 0.009 (n = 6); and TRAM-34, 0.592 ± 0.024 (n = 8). **f** Representative traces and dose-response relationships for whole-cell K^+^ currents upon chemical addition to the extracellular side. Intracellular Ca^2+^ was fixed at 1 μM. IC_50_: propafenone, 56.94 ± 1.04 μM (n = 9); verapamil, 4.60 ± 0.10 μM (n = 8); paxilline, 102.0 ± 1.0 nM (n = 9); TEA, 2.14 ± 0.01 mM (n = 9); and TRAM-34, 247.6 ± 5.6 μM (n = 9). In (**a**), (**b**), (**d**), and (**e**), currents were evoked with a 50-ms pre-pulse at −90 mV pre-pulse for 50 ms, then −90 to 100 mV test pulses for 200 ms with 10-mV increments, and −90 mV post-pulse for 50 ms. Channel activity was calculated as the average current density during the last 50 ms of each test pulse. In (F), currents were evoked by −90 mV pre-pulse for 50 ms, then 100 mV test pulses for 200 ms, then −90 mV post-pulse for 50 ms. Channel activity was calculated as the average current density during the last 50 ms of the test pulse. **g** Viability of 786-O cells subjected to verapamil treatment (0, 10, 30 μM, 48 h) in the presence of high glucose (25 mM), low glucose (1 mM), or the GLUT inhibitor BAY-876 (3 μM). Drug structure (right top). Relative viability for each cell under high/low glucose concentrations (right bottom). **h** Viability of 786-O cells subjected to paxilline treatment (0, 10, 20 μM, 48 h) in the presence of high glucose (25 mM), low glucose (1 mM), or the GLUT inhibitor BAY-876 (3 μM). Drug structure (right top). Relative viability for each cell under high/low glucose concentrations (right bottom). **i** Viability of 786-O cells subjected to TRAM-34 treatment (0, 10, 30 μM, 48 h) in the presence of high glucose (25 mM), low glucose (1 mM), or the GLUT inhibitor BAY-876 (3 μM). Drug structure (right top). Relative viability for each cell under high/low glucose concentrations (right bottom). **j** Viability of 786-O cells subjected to TEA treatment (0, 10, 30 mM, 48 h) in the presence of high glucose (25 mM), low glucose (1 mM), or the GLUT inhibitor BAY-876 (3 μM). Drug structure (right top). Relative viability for each cell under high/low glucose concentrations (right bottom). **k-n** Immunoblotting analysis of cancer-related signaling pathways in 786-O cells treated with verapamil, paxilline, TRAM-34, or TEA (same concentrations as (G)-(J), 8 h), combined with either high (25 mM) or low glucose (1 mM). **o** Total ATP production rates in 786-O cells in high glucose (25 mM); high glucose (25 mM) with propafenone (40 μM); low glucose (0.5 mM); or low glucose (0.5 mM) with propafenone (40 μM). Each condition was treated for 24 h (n = 6). **p** Total ATP production rates in 786-O cells in high glucose (25 mM); high glucose (25 mM) with paxilline (20 μM); low glucose (0.5 mM); or low glucose (0.5 mM) with paxilline (20 μM). Each condition was treated for 24 h (n = 6). **q** Illustration of the LKB1-AMPK-TSC2-mTORC1-S6K pathway that confers the cancer cell-specific cytotoxicity of propafenone/low glucose combination therapy.

Interestingly, we found high expression of large-conductance calcium-activated potassium channels (BK channels), which are a specific subset of K_Ca_ channels (**Fig. 6b, c**). Through patch clamp experiments, we found evidence of BK channel activity in the renal carcinoma cells. In addition, using data from the Cancer Cell Line Encyclopedia (CCLE) database^33^, we observed that several other cancer cell lines used in this study, including A375, 786-O, H358, and BT549, also express various K_Ca_ channels (**Supplementary Fig. 3c-f**).

To further evaluate the involvement of K_Ca_ channels, we conducted patch-clamp analyses to determine whether propafenone inhibits K_Ca_ channel activity. Remarkably, we observed significant inhibition of K_Ca_ channel activity not only with propafenone but also with other antiarrhythmic drugs like verapamil (**Fig. 6d, e**). This activity of verapamil, a CM-SLP^glu^ group 4 drug, further supports the broader therapeutic applicability of other K_Ca_ channel inhibitors. K_Ca_ channel inhibitors, including paxilline, tetraethylammonium (TEA), and Tram-34, demonstrated strong and dose-dependent inhibition of K_Ca_ channel activity (**Fig. 6f**). Importantly, these agents exhibited synergistic cytotoxicity when combined with glucose restriction, mirroring the effects observed with propafenone (**Fig. 6g-j**). To dissect the underlying mechanisms, we next asked if the signaling pathway alterations induced by these drugs were consistent with those elicited by propafenone/low glucose treatment. Interestingly, all tested K_Ca_ channel inhibitors inhibited the mTOR pathway and induced AMPK phosphorylation (**Fig. 6k-n**). These findings suggest a shared mechanism in which K_Ca_ channel inhibition triggers metabolic stress, ultimately modulating the AMPK-mTOR signaling axis to promote cancer cell death.

Following these results, we expanded our investigation to other widely used antiarrhythmic drugs employed in the CM-SLP (**Supplementary Fig. 4a, b**). Amiodarone, a group 3 drug, induced significant K_Ca_ channel inhibition (**Supplementary Fig. 4c, d**) and robust cytotoxicity, along with inhibiting the mTOR pathway, even in the absence of glucose restriction (**Supplementary Fig. 5a, k**).

In contrast, group 1 drugs, as well as procainamide, did not induce sufficient K_Ca_ channel inhibition (**Supplementary Fig. 4c, d**). Neither did these drugs induce significant cell death or inhibit mTOR signaling (**Supplementary Fig. 5b-j, l-t**). These observations not only validate the specificity of K_Ca_ channel inhibition as a therapeutic mechanism but also highlight its potential for repositioning as an anticancer strategy. To confirm that the observed effects were specifically mediated by K_Ca_ channel inhibition, we tested PAP-1, a selective inhibitor of voltage-gated potassium channels. As anticipated, PAP-1 did not affect K_Ca_ channel activity, induce cytotoxicity, or modulate the AMPK-mTOR signaling axis (**Supplementary Fig. 6a-d**), further corroborating the role of K_Ca_ channels in mediating these effects. Given that AMPK is a well-established energy stress sensor, we hypothesized that K_Ca_ channel inhibition, particularly under glucose-restricted conditions, amplifies intracellular energy stress. To test this, we measured ATP production rates under various treatment conditions. Both propafenone treatment and glucose restriction independently reduced ATP production, but their combination resulted in a synergistic and dramatic decline in ATP levels (**Fig. 6o**). We further confirmed this synergistic increase in energy stress by noting that paxilline, a K_Ca_ channel-specific inhibitor, similarly reduced ATP production (**Fig. 6p**). These findings align with previous reports that K_Ca_ channel inhibition reduces calcium flux into mitochondria, impairing ATP synthesis^34^.

Together, our results demonstrate that K_Ca_ channel inhibition, especially when combined with glucose restriction, exacerbates cellular energy stress, leading to cytotoxicity via the LKB1-AMPK-TSC2-mTORC1-S6K signaling axis (**Fig. 6q**). This highlights the therapeutic potential of K_Ca_ channel inhibitors as part of combination treatments targeting cancers with metabolic vulnerabilities, such as glucose dependency. These findings provide a strong rationale for further investigation into the translational application of K_Ca_ channel inhibitors within the CM-SLP framework, paving the way for innovative cancer therapies tailored to exploit tumor-specific metabolic stress conditions.

### Biguanide/low glucose impairs metabolic plasticity and Hippo-YAP signaling in cancer cells

Next, we conducted experiments to explore the molecular mechanism of another top-scoring CM-SLP^glu^ candidate, phenformin, a biguanide, within the CM-SLP group 4. The aim was to validate the anti-cancer efficacy of biguanide/low glucose treatment and identify alternative targets that could serve as substitutes for hypoglycemia.

To further validate the cancer cell-specific cytotoxicity of biguanide/low glucose, we subjected melanoma, NSCLC, and breast cancer cells alongside their normal counterparts to daily cycles of combined treatment. Although each cell line showed differential sensitivity, metformin elicited significant cytotoxicity under glucose-restricted conditions in all cancer cell lines, but not in non-cancer cells (**Fig. 7a**). To understand the cooperative effect of biguanide and low glucose on cellular energy status, we measured the ATP production rate derived via either glycolysis or mitochondrial oxidative phosphorylation (OXPHOS) (**Fig. 7b**). It is important to note that neither biguanide nor glucose restriction alone impaired intracellular ATP production. Metformin treatment shifted ATP production toward glycolysis, whereas glucose restriction shifted ATP production toward OXPHOS, demonstrating the metabolic plasticity of cancer cells. These results show that combined treatment is necessary to induce a significant reduction in ATP production and total cellular ATP levels (**Fig. 7c**).

**Fig. 7.**
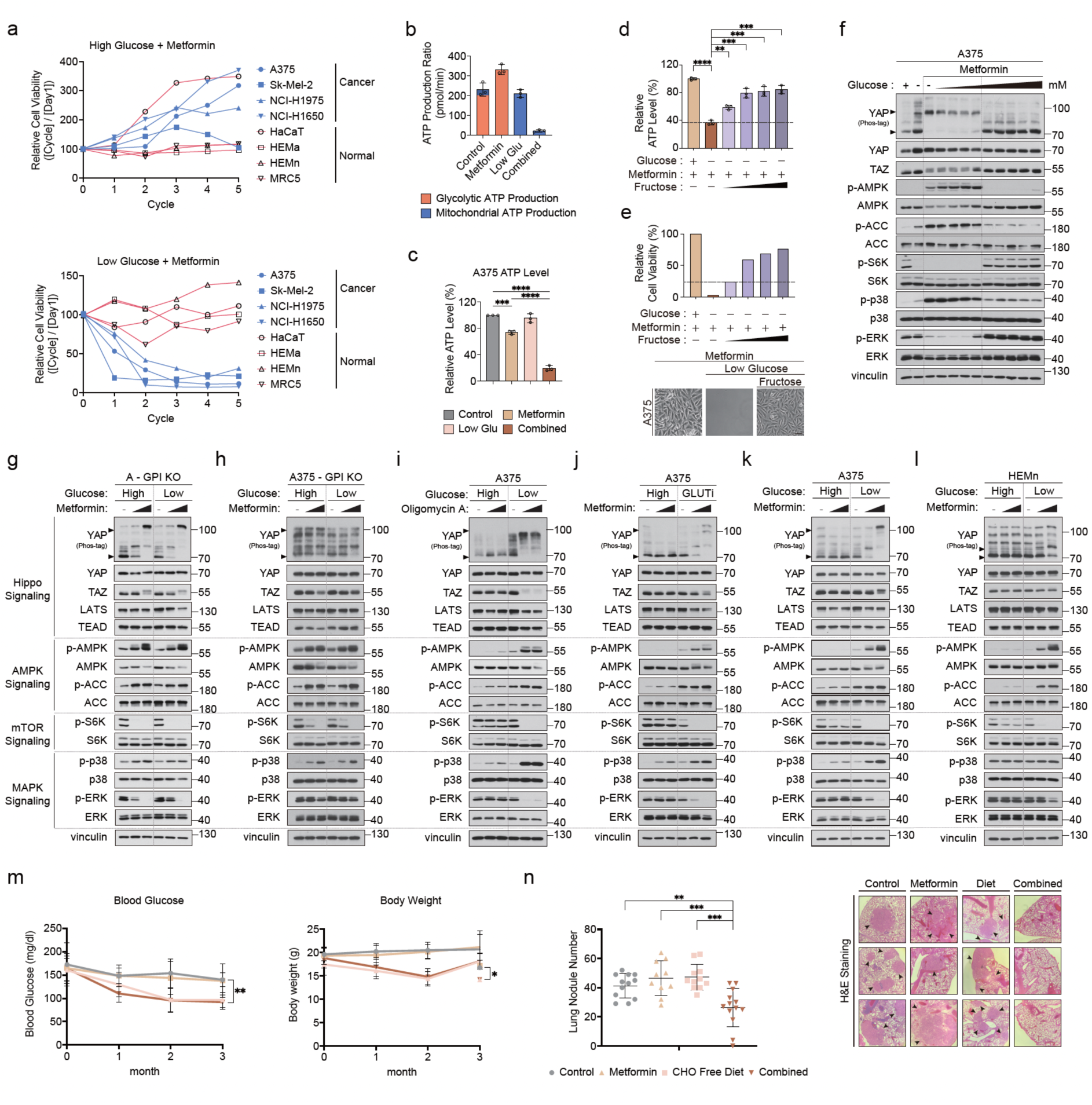
Biguanide/low glucose impairs metabolic plasticity and Hippo-YAP signaling in cancer cells. **a** Comparing the viability of cancer cells and normal cells subjected to daily cycles (24 h) of metformin/high glucose (top) or metformin/low glucose (bottom) treatments. Metformin, 10 mM; high glucose, 25 mM; low glucose, 1 mM. Each data point in the graph was calculated using a two-step normalization process. First, relative cell viability for each cycle was calculated as follows: relative cell viability = (cell viability of glucose + metformin group)/(cell viability of glucose control group at the same time point) × 100. Next, these values were normalized to cycle 0 as follows: normalized relative cell viability = (relative cell viability at cycle X)/(relative cell viability at cycle 0) × 100. **b** Measurements of real-time ATP production (pmol/min) in A375 cells subjected to either metformin alone, low glucose alone, or in the two in combination. **c** Measurements of relative intracellular ATP levels in A375 cells treated as in (B). **, p < 0.01; ****, p < 0.0001. Statistics were analyzed via one-way ANOVA with Bonferroni corrections for multiple comparisons. **d, e** Viability and intracellular ATP levels measured in A375 cells after fructose supplementation. Addition of fructose (25–200 mM) rescued metformin/low glucose-induced ATP deprivation and cancer cell death in a dose-dependent manner. **, p < 0.01; ***, p < 0.001; ****, p < 0.0001. Statistics were analyzed via one-way ANOVA with Bonferroni corrections for multiple comparisons. **f** Immunoblotting analysis of A375 cells treated with various glucose concentrations (0.5, 1, 2, 2.5, 5, 10, 15, 20, and 25 mM, 11 h) combined with metformin (10 mM, 8 h). Glucose concentrations lower than 2.5 mM synergized with metformin to disrupt cell signaling. The arrowheads indicate YAP phosphorylation status. **g** Immunoblotting analysis of glucose-6-phosphate isomerase (GPI) knockout HEK293 cells treated with metformin (0, 3, 10 mM, 8 h) in high (25 mM) or low glucose (1 mM, 11 h). **h** Immunoblotting analysis of glucose-6-phosphate isomerase (GPI) knockout A375 cells treated with metformin (0, 3, 10 mM, 8 h) in high (25 mM) or low glucose (1 mM, 11 h). **i** Immunoblotting analysis of signal pathway perturbations in A375 cells treated with oligomycin A (0, 1, 5 μM, 8 h) combined with high (25 mM) or low glucose (1 mM, 11 h). **j** Immunoblotting analysis of A375 cells treated with metformin (0, 3, 10 mM, 8 h) combined with high glucose (25 mM) or the GLUT inhibitor BAY-876 (10 μM, 11 h). **k** Immunoblotting analysis of A375 cells treated with metformin (0, 3, 10 mM, 8 h) in high (25 mM) or low glucose (1 mM, 11 h). **l** Immunoblotting analysis of HEMn cells treated with metformin (0, 3, 10 mM, 8 h) in high (25 mM) or low glucose (1 mM, 11 h). **m** Measurements of blood glucose levels (mg/dl) and body weights (g) from each group for 3 months. *, p < 0.05; ****, p < 0.001. **n** Number of lung tumor nodules counted from each experimental group. Lung nodules were counted using microscopy in representative images of H&E-stained lungs from each mouse group. Black arrows indicate tumor nodules. **, p < 0.01.

To determine whether restoring ATP production can rescue biguanide/low glucose-induced cell death, we supplied cells with fructose, which bypasses the pentose phosphate pathway (PPP) and the hexosamine pathway to directly feed the glycolytic pathway. This provides cells an alternative way to produce ATP even under conditions that impair both glucose uptake and OXPHOS (**Fig. 7d**). We found that fructose restored ATP production and increased cell survival in a dose-dependent manner, suggesting biguanide in combination with glucose restriction promotes cell death via induction of energy crisis (**Fig 7e**). We next examined the dysregulation of energy stress signaling via the Hippo, AMPK, mTOR, and MAPK signaling pathways to determine whether physiologic glucose levels cooperate with biguanides to induce such signaling perturbations in cancer cells. We found that reduction of glucose levels below 5 mM was sufficient to disrupt key signaling pathways, suggesting that the combination of biguanide and low glucose affects cancers at physiologically relevant hypoglycemic conditions (**Fig. 7f**). Mechanistically, depletion of glucose-6-phosphate isomerase (GPI, the second enzyme in the glycolytic pathway), which impairs downstream glycolytic ATP production, elicited biguanide-induced signaling perturbations even in high glucose (**Fig. 7g, h**). Moreover, the substitution of metformin treatment or glucose restriction with either the mitochondrial ATP synthase inhibitor oligomycin or a GLUT inhibitor evoked identical signaling alterations (**Fig. 7i, j**). These results confirm that biguanide and low glucose cooperate to target metabolic plasticity between glycolysis and OXPHOS.

Next, we examined cancer-related signaling pathways to identify the mechanism by which the combination of biguanide and hypoglycemia differentially affects the viability of cancer cells versus normal cells. Metformin alone failed to trigger changes in signaling pathways in melanoma cells. Under low glucose, however, metformin evoked aberrant cellular responses, resulting in a marked disruption of multiple signaling pathways. These included YAP inhibition, AMPK activation, mTOR inhibition, ERK inhibition, and p38 activation, indicating that the combined treatment induces energy depletion, cellular stress, and growth suppression (**Fig. 7k**). The combined treatment also affected these same pathways in BRAF inhibitor-resistant melanoma cells, as well as NSCLC cells, breast cancer cells, gastric cancer cells, and head and neck cancer cells (**Supplementary Fig. 7a-f**). We next asked whether differential signaling perturbations underlie the contrasting vulnerability of normal counterpart cell lines. In normal melanocytes, lung fibroblasts, and non-malignant breast epithelial cells, metformin/low glucose recapitulated the pathway perturbations observed in cancer cells, except for Hippo pathway-induced YAP/TAZ inhibition (**Fig. 7l, Supplementary Fig. 7g-i**). YAP and TAZ are oncogenic transcriptional coactivators and effectors of the Hippo pathway that, in response to various stress and growth stimuli, play critical roles in tumor progression and regeneration^35–37^. We observed YAP/TAZ phosphorylation and reduced YAP/TAZ stability exclusively in cancer cells, suggesting differential regulation of the Hippo pathway confers cancer cell-specific metabolic vulnerability and cytotoxicity via biguanide/low glucose treatment^38–40^.

To confirm the anticancer efficacy of biguanides under hypoglycemic conditions, we treated K-Ras^LA2^ lung cancer model with the combination of metformin and either a normal or carbohydrate-free diet. Notably, mice exposed to a carbohydrate-free diet alone showed reduced blood glucose and body weight, whereas neither metformin alone nor metformin combined with a carbohydrate-free diet induced any further reduction in either parameter (**Fig. 7m**). These results are consistent with previous reports indicating metformin lowers blood glucose levels in type 2 diabetic patients but not in subjects with normal glucose levels^41^. Importantly, we found the anti-cancer effect of metformin was highly dependent on blood glucose level. Metformin had a minor effect on lung tumor growth in mice fed with a normal diet, as did the carbohydrate-free diet alone. The combination of metformin and carbohydrate-free diet, however, induced hypoglycemia and dramatically synergized to impair tumor growth (**Fig. 7n**). Our CM-SLP^glu^ results suggest the antidiabetic biguanides, including phenformin and metformin, are highly sensitive to glucose restriction. Not only does this resolve the controversial anticancer effect of the biguanides^42^, it also strengthens previous reports by demonstrating that, among the 1,813 FDA-approved drugs, the biguanides are one of the most prominent drugs that synergize with the hypoglycemic condition^43–45^

### Biguanide/low glucose targets YAP-driven cancer cells via the ATP-FAK-RhoA-LATS axis

Next, we aimed to identify the molecular mechanism by which the combination of biguanide and glucose restriction links ATP depletion to YAP inhibition. Morphologically, cancer cells exhibited cell shrinkage and disturbed focal adhesions (FA), suggesting that cellular energy stress-induced cytotoxicity is mediated at least in part through the disruption of integrin-FAK signaling, FA assembly, and F-actin remodeling in cancer cells (**Fig. 8a, b**). By restoring cellular ATP levels with fructose, we were able to rescue FAK activation and LATS inhibition followed by dephosphorylation and activation of YAP/TAZ (**Fig. 8c**). We next tested the involvement of RhoA and the Hippo components LATS1 and 2 to determine whether regulation of the RhoA-Hippo pathway mediates focal adhesion disassembly and YAP inhibition. Notably, we found ectopic expression of constitutively active RhoA (**Fig. 8d**) and depletion of LATS1/2 (**Fig. 8e**) both dramatically blocked YAP/TAZ phosphorylation induced by biguanide/low glucose treatment. Although the LKB1-AMPK axis is a well-established energy sensor and upstream suppressor of YAP/TAZ, depletion of AMPK failed to rescue YAP/TAZ inhibition (**Fig. 8f**). Hippo pathway-induced YAP phosphorylation subsequently led to YAP cytoplasmic translocation (**Fig. 8g**) and disruption of YAP-TEAD interaction, resulting in reduced expression of TEAD target genes that derive cancer progression (**Fig. 8h-j**). These results indicate that the blockade of metabolic plasticity that evokes cancer cell-specific cytotoxicity occurs via the FAK-RhoA-LATS-YAP/TAZ-TEAD signaling axis (**Fig. 8k**).

**Fig. 8.**
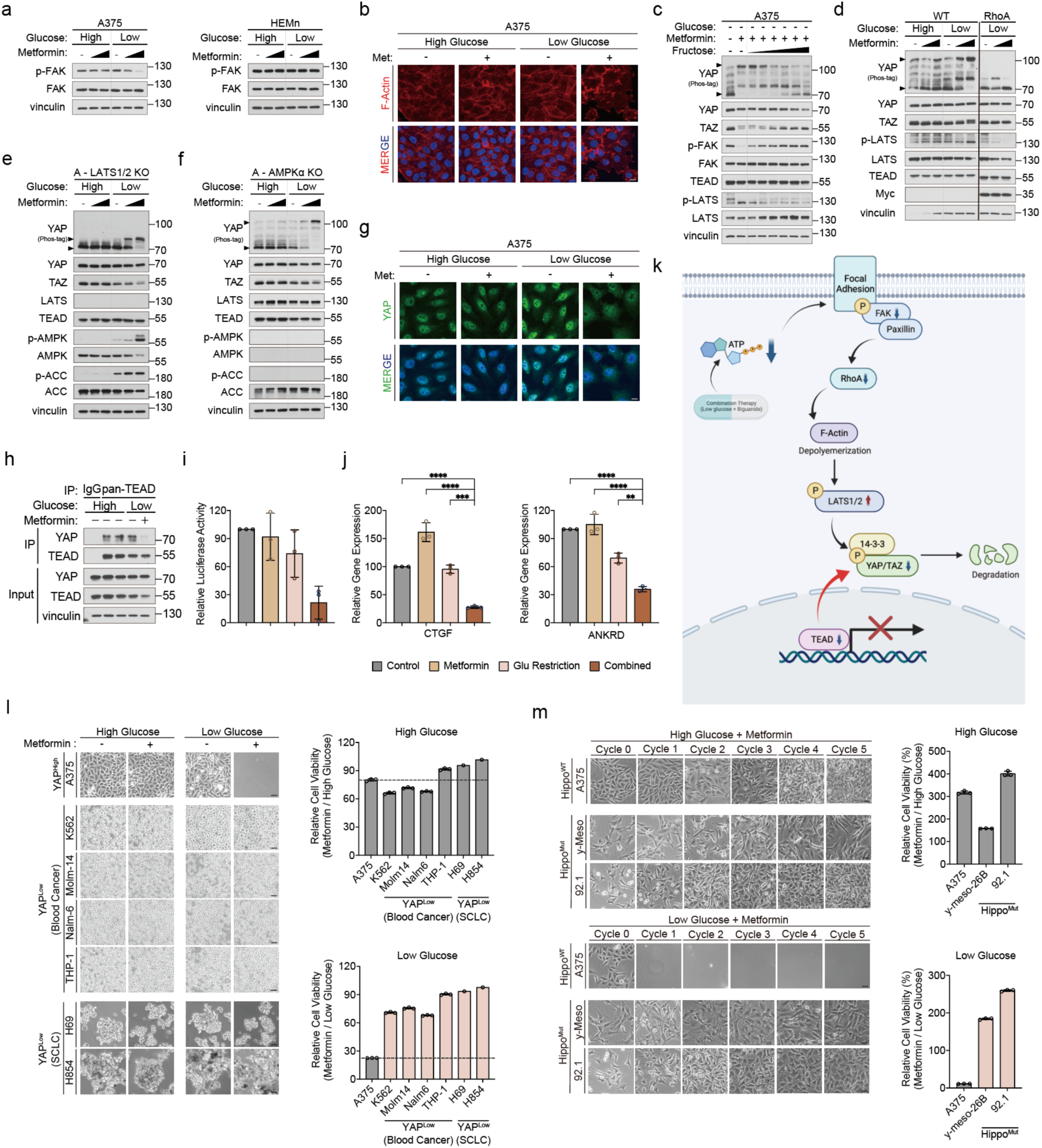
Biguanide/low glucose targets YAP-driven cancer cells via the ATP-FAK-RhoA-LATS axis. **a** Immunoblotting analysis comparing FAK phosphorylation levels in A375 and HEMn cells after treatment with metformin (0, 3, 10 mM, 8 h) in high (25 mM) or low glucose (1 mM, 11 h). **b** Immunofluorescence images of F-actin depolymerization in A375 cells treated with metformin (0, 3, 10 mM, 8 h) in high (25 mM) or low glucose (1 mM, 11 h). Red, phalloidin; blue, DAPI. **c** Immunoblotting analysis of lysates derived from A375 cells treated with metformin (0, 3, 10 mM, 8 h) in high (25 mM) or low glucose (1 mM, 11 h) supplemented with fructose (10, 25, 50, 100, 150, and 200 mM) as in Figure 7D. **d** Immunoblotting analysis of YAP and LATS dephosphorylation induced via ectopic RhoA expression in HEK293 cells treated with metformin (0, 3, 10 mM, 8 h) in high (25 mM) or low glucose (1 mM, 11 h). **e, f** Immunoblotting analysis of YAP phosphorylation status in LATS1/2 knockout or AMPKα knockout HEK293A cells treated with metformin (0, 3, 10 mM, 8 h) in high (25 mM) or low glucose (1 mM, 11 h). **g** Immunofluorescence analysis of YAP cytoplasmic translocation in A375 cells treated with metformin (0, 3, 10 mM, 8 h) in high (25 mM) or low glucose (1 mM, 11 h). Green, YAP; blue, DAPI. **h** Immunoprecipitation analysis of the interaction between YAP and TEAD in the cell lysates used in (G). **i** TEAD luciferase reporter activity from the 8xGTIIC plasmid in HEK293A cells subjected to metformin/low glucose treatment. Y-axis, relative luciferase units (RLU). n = 3. *, p < 0.05, **, p < 0.01. Statistics were analyzed via t-test. **j** Quantitative real-time PCR analysis of the TEAD target genes CTGF and ANKRD1 in HEK293A cells subjected to metformin/low glucose treatment. n = 3. *, p < 0.05; **, p < 0.01, ***, p < 0.001; ****, p < 0.0001. Statistics were analyzed via one-way ANOVA with Bonferroni corrections for multiple comparisons. **k** Illustration of the ATP-FAK-RhoA-LATS-YAP/TAZ-TEAD pathway, which confers the cancer cell-specific cytotoxicity of biguanide/low glucose combination therapy. **l** Viability of YAP^high^ and YAP^low^ cancer cells treated with metformin in high (25 mM) or low glucose (1 mM). YAP^high^ cell line, A375; YAP^low^ cell lines: the K562, Molm-14, Nalm-6, and THP-1 blood cancer cell lines, as well as the NCI-H69 and NCI-H854 small cell lung cancer cell lines. **m** Viability of Hippo^OFF^/YAP^ON^ cancer cells subjected to metformin in high (25 mM) or low (1 mM) glucose media. Hippo^OFF^/YAP^ON^ cells are resistant to biguanide/low glucose treatment. The Hippo^OFF^/YAP^ON^ cancer cell lines included the y-meso-26B mesothelioma and 92.1 uveal melanoma cell lines.

Because YAP and TAZ mediate biguanide/low glucose-induced cancer cell death, it is conceivable that the combined treatment induces minimal cytotoxicity in YAP/TAZ-deficient (YAP^low^) cancer cells. Although the pro-tumorigenic role of YAP/TAZ is well-established in various solid tumors, some hematological malignancies and SCLC cells hardly express YAP/TAZ^46,47^. In contrast to our results with YAP-expressing cancer cells (YAP^high^), the combined treatment hardly elicited any cell death in YAP-deficient SCLC and blood cancer cell lines, indicating that YAP/TAZ expression is an indicator of cell vulnerability to the biguanide/low glucose combination therapy (**Fig. 8l**). Because LATS1/2-mediated Hippo signaling is required to relay energy stress signals to YAP/TAZ inhibition, we next asked whether Hippo-inactivated cancer cells expressing constitutively active YAP (Hippo^OFF^/YAP^ON^) exhibit resistance to combined therapy. In comparison to Hippo wild type (WT) cells, Hippo^OFF^/YAP^ON^ cancer cells, such as 92.1 uveal melanoma cells and y-MESO-26B mesothelioma cells, which express constitutively active YAP, showed marked resistance to comined biguanide/low glucose treatment (**Fig. 8m**). These results demonstrate that cancer cell-specific Hippo signaling perturbations mediate the anticancer effect of biguanide/low glucose, thus rendering Hippo^OFF^/YAP^ON^ cancer subtypes highly resistant to such combined treatment. Therefore, we aim to overcome these limitations of Hippo^OFF^/YAP^ON^ cancer cells and dietary interventions by utilizing YAP-TEAD inhibitors that can bypass Hippo kinases via direct targeting of TEAD.

### Redirecting metabolic intervention toward YAP-TEAD targeted therapies improves the clinical applicability of CM-SLP

Because it remains challenging to apply hypo-nutrient metabolic interventions in human patients, we explored alternative means to exploit targeted therapies suggested by CM-SLP^glu^ that would improve their clinical applicability. Since Hippo components and YAP status are critical determinants of cancer cell susceptibility to biguanide/hypoglycemia, we developed a small molecule inhibitor of the YAP-TEAD complex to circumvent dietary interventions and further target the non-responsive Hippo^OFF^/YAP^ON^ cancer cells. We performed a computer-based drug screen to identify novel inhibitors of YAP-TEAD protein-protein interactions (PPI) that target TEAD interface 3 and dissociate YAP/TAZ from TEAD (**Supplementary Fig. 8a**). After generating pharmacophore models for the design of inhibitors that bind interface 3 of the YAP–TEAD PPI, we performed a virtual screen of 7 million small molecules to identify BMY-123 (**Supplementary Fig. 8b**)^48^. We used the semi-parameterized fragment molecular orbital (FMO) method to analyze essential interaction energies within the BMY-123/TEAD complex. We identified E368, E393, and D249 in TEAD1 as residues that are particularly important for its interaction with BMY-123 (**Supplementary Fig. 8c**). We also found that electrostatic (ES) and hydrophobic (DI) interaction energies are critical for the BMY-123 and TEAD1 interaction (**Supplementary Fig. 8d**). Surface plasmon resonance analysis indicated that BMY-123 binds directly to TEAD1 with a KD value of 24.1 mM (**Supplementary Fig. 8e**).

Because inhibition of YAP-TEAD signaling mediates the cytotoxicity of biguanide/hypoglycemia combined treatment, we next asked whether BMY-123 could act as a substitute for hypoglycemia in synergizing with biguanides. Combination of biguanide/BMY-123 treatment potently dissociated YAP/TAZ from TEAD (**Supplementary Fig. 8f**). Although BMY-123 alone was sufficient to suppress TEAD reporter activity and target gene expression, the combination of phenformin and BMY-123 significantly enhanced its inhibitory effect on TEAD (**Supplementary Fig. 8g**). Consistent with these results, the combination of phenformin and BMY-123 enhanced cytotoxicity beyond that induced by BMY-123 alone, suggesting BMY-123 can improve the clinical feasibility of biguanide/hypoglycemia treatment by replacing the hypoglycemic condition (**Supplementary Fig. 8h**). To take further advantage of the biguanide/BMY-123 combination, we asked whether it could overcome the limitation of biguanide/hypoglycemia combination that failed to trigger cytotoxicity in Hippo^OFF^/YAP^ON^ cancer cells. In comparison to BMY-123 alone, phenformin/BMY-123 treatment markedly enhanced cytotoxicity in A375 melanoma cells and in the Hippo^OFF^/YAP^ON^ 92.1 uveal melanoma cells that were insensitive to biguanide/low glucose treatment (**Fig. 8m, Supplementary Fig. 8h**). Notably, Hippo^OFF^/YAP^ON^ cells showed an even lower IC_50_ to combined treatment, possibly due their YAP-dependency. In addition, the combination of biguanide and BMY-123 markedly improved the IC_50_ for cancer cells compared to their normal counterparts (**Supplementary Fig. 8h**).

To verify the *in vivo* efficacy of biguanide/BMY-123 treatment, we treated K-Ras^LA2^ mice daily with metformin and BMY-123, both alone and in combination. Compared to metformin or BMY-123 alone, combined treatment with metformin and BMY-123 showed a synergistic effect and significantly enhanced anticancer efficacy *in vivo*. Thus, metformin/BMY-123 recapitulated the therapeutic effect of metformin/hypoglycemia even in the absence of dietary intervention (**Supplementary Fig. 8i**). Additionally, we found BMY-123 exhibited better synergistic effects with phenformin compared to the recently reported TEAD inhibitor flufenamic acid (**Supplementary Fig. 8j, k**)^49^. Together, these results highlight the critical role of the Hippo pathway in mediating the synergistic anticancer effect of biguanides and hypoglycemia *in vitro* and *in vivo*. Thus, we have provided the rationale for expanding the use of CM-SLP to identify the critical determinants of metabolic drug vulnerabilies and further exploit undescribed combinatorial targeted therapies for improving clinical applicability.

## Discussion

In this study, we introduced CM-SLP, a metabolite-dependent high-throughput screening platform. CM-SLP was designed to guide the repurposing of a panel of FDA-approved non-oncology and oncology drugs based on their unexpected cytotoxicity and metabolic vulnerability in conditions of altered nutrient availability. Our results expand the current concept of precision medicine toward individualized medication guidelines that depend on each cancer patient’s metabolic state. Growing evidence demonstrates the dynamic interplay between cancer metabolism and drug efficacy, indicating that metabolic status, either system-wide or within the tumor microenvironment, can significantly affect a patient’s response to various drugs^4,6,50^. Hyper-nutrient states, such as those occurring in the presence of metabolic syndromes, are considered major risk factors of cancer incidence, whereas hypo-nutrient states obtained via dietary interventions, such as intermittent fasting or calorie restriction, can potentiate anticancer drug efficacy^9,51,52^. Hence, CM-SLP provides a comprehensive resource for the analysis and prediction of an individual cancer patient’s drug response according to his or her metabolic status. Remarkably, CM-SLP identifies numerous FDA-approved non-oncology and oncology drugs that have not been previously reported to affect cancer viability in the context of altered nutrient availability. Moreover, in addition to guiding the treatment of cancer patients with repurposed non-oncology drugs that match their associated metabolic conditions, CM-SLP also suggested the possibility that metabolic status may explain unwanted cytotoxicity from non-oncology drugs in non-cancer patients.

Previous research has shown that nutrient levels in the blood plasma and microenvironmental tumor interstitial fluid (TIF) dictate nutrient availability in cancer cells^8^. Both cell-intrinsic and extrinsic factors, such as tumor type, epigenetic state, anatomic location, tumor microenvironment, and other cellular components can impact cancer cell metabolism. Importantly, the metabolic composition of TIF differs from that of plasma. Moreover, because systemic metabolic changes affect the metabolite composition of TIF, it is likely that metabolic syndromes or dietary interventions influence tumor nutrient availability by altering the levels delivered from the circulation, allowing them to synergize with group 2 and 4 CM-SLP candidates. Further research is warranted on how cancer cells respond to CM-SLP-derived drugs with differential sensitivity depending on systemic and local metabolite concentrations and how non-cancer cells, such as immune cells and fibroblasts, within the tumor microenvironment influence this differential sensitivity. Moreover, we also consider it important to find standardized ways for individual patients to assess complex metabolic characteristics to guide the administration of CM-SLP drugs.

CM-SLP identified glucose as the major metabolite conferring differential drug responses with the highest selectivity toward cancer cells compared to their normal counterparts. Although previous studies indicate that cancer growth depends on several different metabolites, we found it intriguing that, glucose, rather than glutamine or fatty acids, contributed the most to endowing non-oncology drugs with marked cytotoxicity in various cancer types. It is noteworthy that, distinct cancer types may respond with different CM-SLP scores for each condition. Moreover, while most studies have examined how reducing glucose levels potentiates drug responses, we initiated a follow-up study on the group 2 drug fenbendazole, which exhibited unexpected cytotoxicity in elevated glucose conditions. We found that, in the presence of high glucose, fenbendazole induced YAP protein band shift, indicating disrupted YAP signaling. It is important to note that glucose concentrations above 10 mM (180 mg/ dL) were required for fenbendazole to induce significant cytotoxicity, suggesting cancer patients with hyperglycemia could benefit from fenbendazole treatment. To our knowledge, this is the first evidence demonstrating that high glucose can potentiate the cytotoxicity of non-oncology drugs. Consistent with these results, hyperglycemia potentiates chemotherapy responses in pancreatic cancer patients and mouse models^53^. We also confirmed that drugs selected from both glucose and glutamine panels exhibited changes in anticancer efficacy based on metabolite concentrations (**Supplementary Fig. 2**). Thus, we expect further research on the hyper/hypo-nutrient-associated cytotoxicity of group 2 and 4 CM-SLP drugs will offer new approaches for the repurposing of various FDA-approved drugs.

On the other hand, although hyper-nutrient conditions that synergize with group 2 CM-SLP drugs are clinically relevant and readily applicable, hypo-nutrient conditions that work together with group 4 CM-SLP candidates are challenging to achieve in cancer patients^52^. It is therefore critical to identify dysregulated signaling pathways that are relevant to cancer cell-specific cytotoxicity and substitute dietary interventions with a more readily applicable targeted therapy. Surprisingly, group 4 CM-SLP^glu^ scores indicate that, among FDA-approved drugs, glucose restriction confers significant cytotoxicity to propafenone and biguanides. We found that dual inhibition of glycolysis and calcium flux, achieved via combination of glucose restriction with propafenone (K_Ca_ channel blocker) induced energy stress in cancer cells. Clarifying the mechanisms of hypo-nutrient-sensitive CM-SLP drugs mediated via AMPK-mTOR inhibition, we found that targeted treatment with temsirolimus, which hinders mTOR complex formation, could partially replace the hypoglycemia requirement. Additionally, we and others found that dual inhibition of glycolysis and OXPHOS, achieved via combination of glucose restriction and biguanides, was required to suppress metabolic plasticity and ATP production in cancer cells^44,45,54^. Further clarifying the mechanisms of hypo-nutrient-sensitive CM-SLP drugs mediated via YAP-TEAD inhibition, we found that targeted treatment with BMY-123, which disrupts the YAP-TEAD complex, could partially replace the hypoglycemic requirement. Although Hippo^OFF^/YAP^ON^ status renders cancer cells insensitive to the biguanide/low glucose combination, the alternative biguanide/BMY-123 combination allowed us to overcome this limitation. In addition to these combination therapies, we validated the cytotoxicity and altered signaling pathways of other top group 4 CM-SLP^glu^ combinations, including valdecoxib/hypoglycemia, verapamil/hypoglycemia, and nilutamide/hypoglycemia. According to our results, these high-scoring CM-SLP combination therapies disrupted similar cancer signaling pathways, in part via their ability to target mitochondrial activity. Further identification of the signaling mechanisms and mediators of group 4 CM-SLP candidates will provide the rationale for novel combinations of metabolic interventions that will facilitate drug repurposing and clinical application of CM-SLP.

Here, we performed a large-scale screen to identify anticancer uses of non-oncology FDA-approved drugs, focusing on those with differential cytotoxicity in hyper- and hypo-metabolite conditions. The appeal of drug repurposing based on CM-SLP data is the rapid clinical translation of findings with drugs that have already proven safe in humans. The non-oncology drugs used in our screening showed low cytotoxicity when used alone even at higher concentrations. This suggests their safety profile is robust, as even higher doses fail to induce significant cell death in non-cancerous settings. Given the large number of drug repurposing candidates with unexpected metabolite dependencies that emerged in this initial screen, we aim to expand CM-SLP approach toward various cancer types, metabolites, and mouse models. In addition to the immediate repurposing of existing drugs for anticancer indications, the CM-SLP database introduces a framework that can guide the way to a treatment modality we refer to as oncometabolic precision medicine. The idea is to provide medication guidelines for individual patents based on metabolite concentrations in their blood or tumor microenvironments. This will help diversify anticancer treatment modalities and maximize the safety and efficacy of such treatments. Our CM-SLP results further suggest that, during preclinical and clinical trials, drug efficacy and cytotoxicity should be examined under key representative metabolic conditions to improve clinical decision making.

---

**Supplementary Fig. 1:**
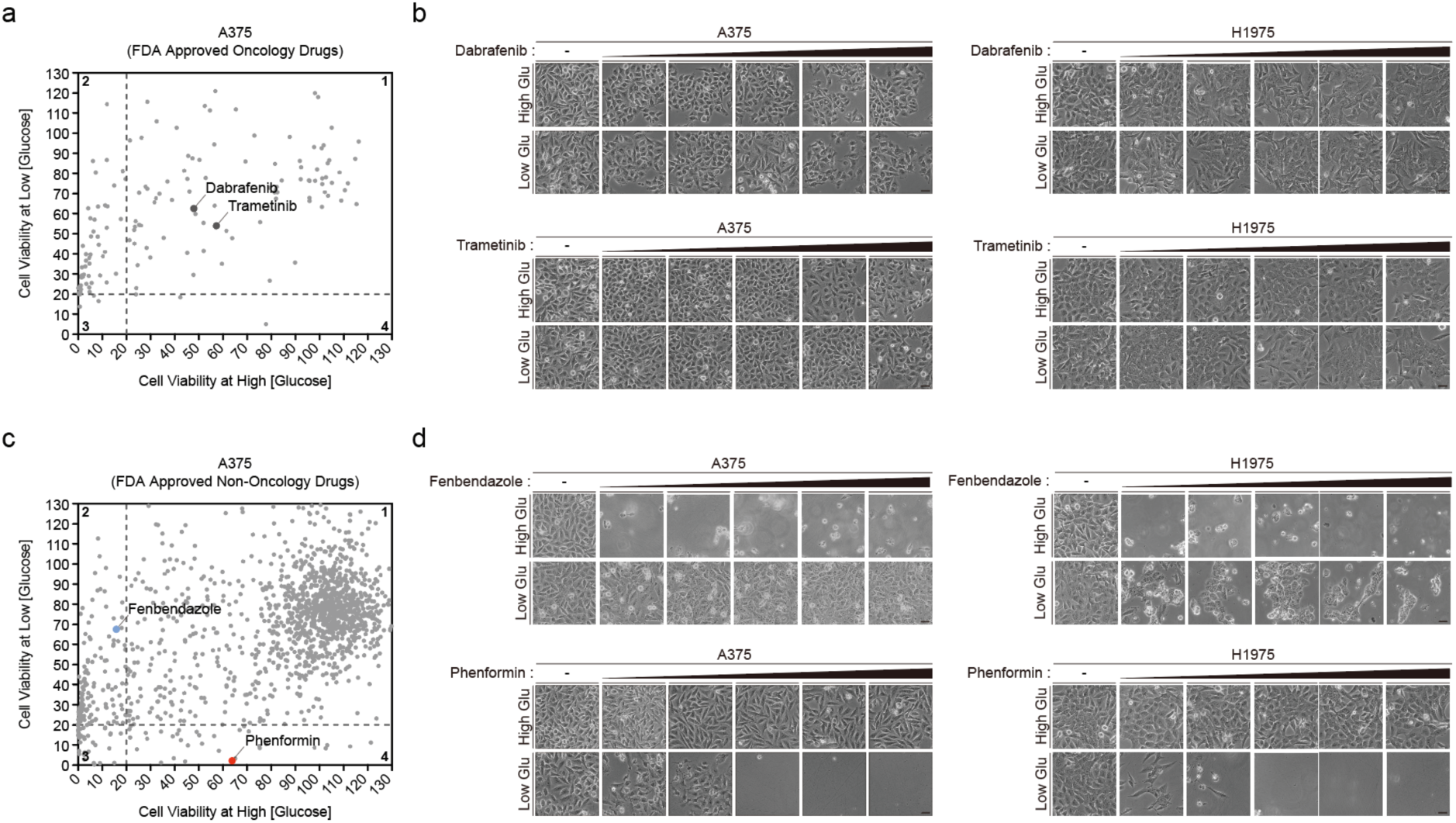
Glucose availability affects the anticancer potential of non-oncology drugs in vivo. **a** Dot plot of FDA-approved oncology drugs from the CM-SLP^glu^ screen showing relative cell viability under high glucose (x-axis) versus low glucose (y-axis). Dabrafenib and trametinib (group 1) are marked. **b** Viability of A375 and H1975 cells subjected to the indicated concentrations of the group 1 drugs dabrafenib or trametinib (0, 20, 40, 60, 80, 100 μM, 24 h) in the presence of high (25 mM) or low glucose (1 mM, 27 h). **c** Dot plot of FDA-approved non-oncology drugs from the CM-SLP^glu^ screen showing relative cell viability under high glucose (x-axis) versus low glucose (y-axis). Fenbendazole (group 2) and phenformin (group 4) are marked. **d** Viability of A375 and H1975 cells subjected to the indicated concentrations of either the group 2 drug fenbendazole or the group 4 drug phenformin (0, 20, 40, 60, 80, 100 μM, 24 h) in the presence of high (25 mM) or low glucose (1 mM, 27 h).

**Supplementary Fig. 2:**
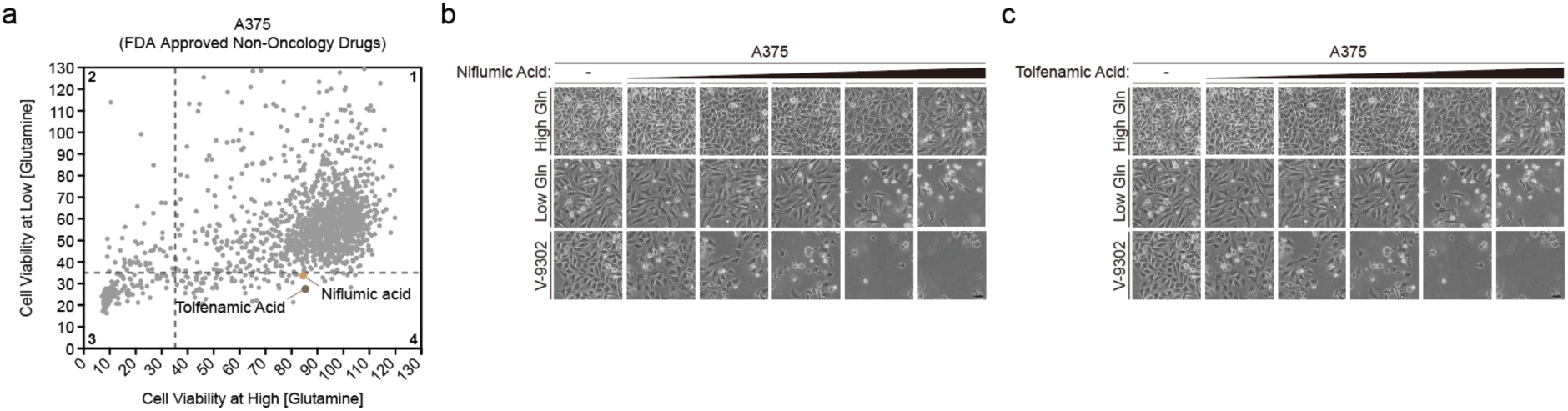
Glutamine availability affects the anticancer potential of non-oncology drugs. **a** Dot plot of FDA-approved non-oncology drugs from the CM-SLP^gln^ screen showing relative cell viability under high glucose (x-axis) versus low glucose (y-axis). Niflumic acid and tolfenamic acid (group 4) are marked. **b** Viability of A375 cells subjected to the indicated doses of the group 4 drug niflumic acid (0, 100, 150, 200, 300, 500 μM, 24 h) in the presence of high glutamine (2 mM), low glutamine (0 mM, 27 h), or the ASCT2 inhibitor V-9302 (30 μM, 27 h). **c** Viability of A375 cells subjected to the indicated doses of the group 4 drug tolfenamic acid (0, 100, 150, 200, 300, 500 μM, 24 h) in the presence of high glutamine (2 mM), low glutamine (0 mM, 27 h), or the ASCT2 inhibitor V-9302 (30 μM).

**Supplementary Fig. 3:**
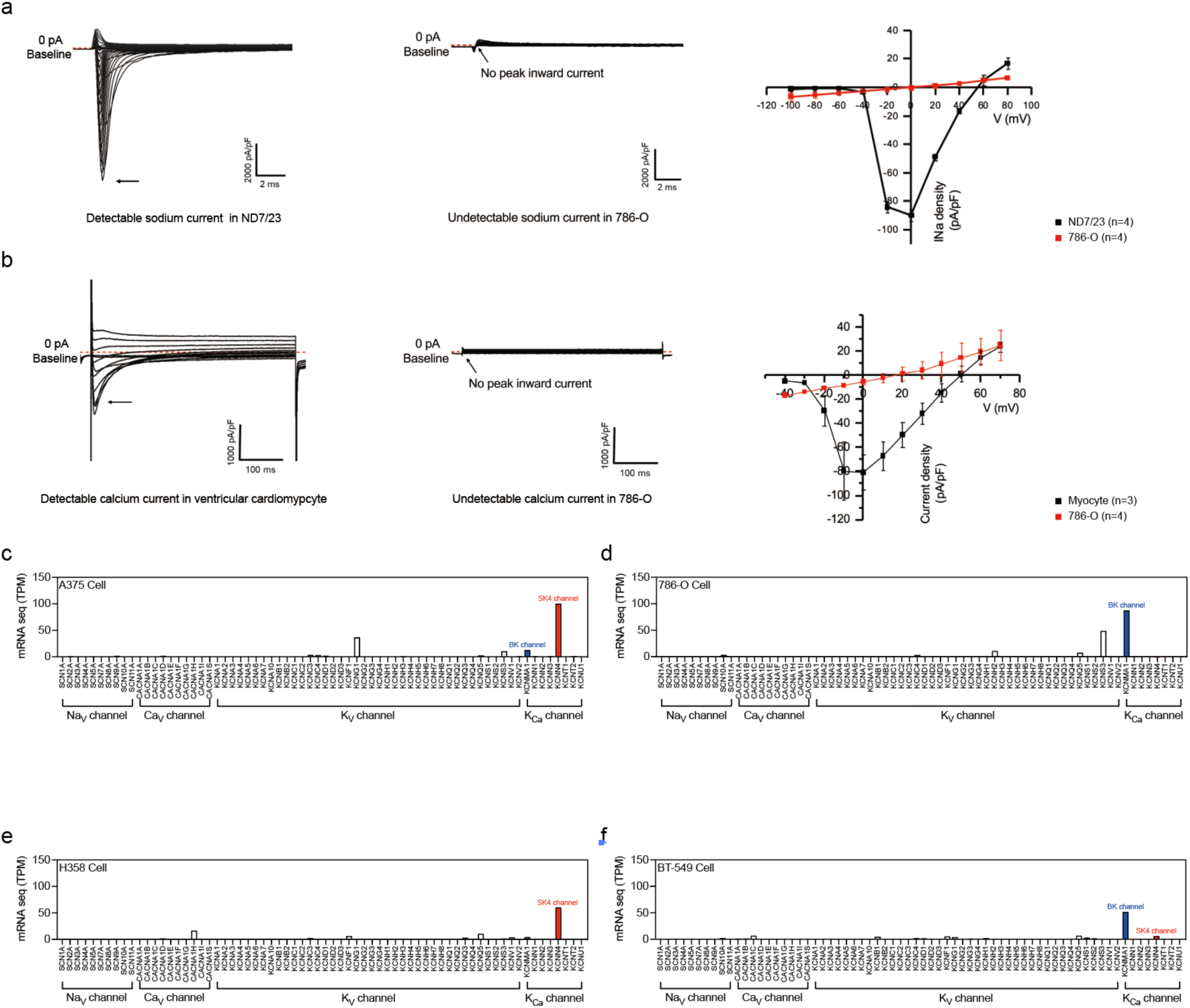
Anticancer effects of propafenone are not mediated via its canonical target. **a** Voltage-activated Na^+^ (Na_V_) currents in ND7/23 and 786-O cells. **b** Voltage-activated Ca^2+^ (Ca_V_) currents in ventricular cardiomyocytes and 786-O cells. **c-f** Transcripts per million (TPM) values for voltage-activated Na^+^ channels, voltage-activated Ca^2+^ channels, voltage-activated K^+^ channels, and Ca^2+^-activated K^+^ channels obtained from the Cancer Cell Line Encyclopedia (CCLE) database (Ref). (C) A375 cells, (D) 786-O cells, (E) H358 cells, and (F) BT-549 cells.

**Supplementary Fig. 4:**
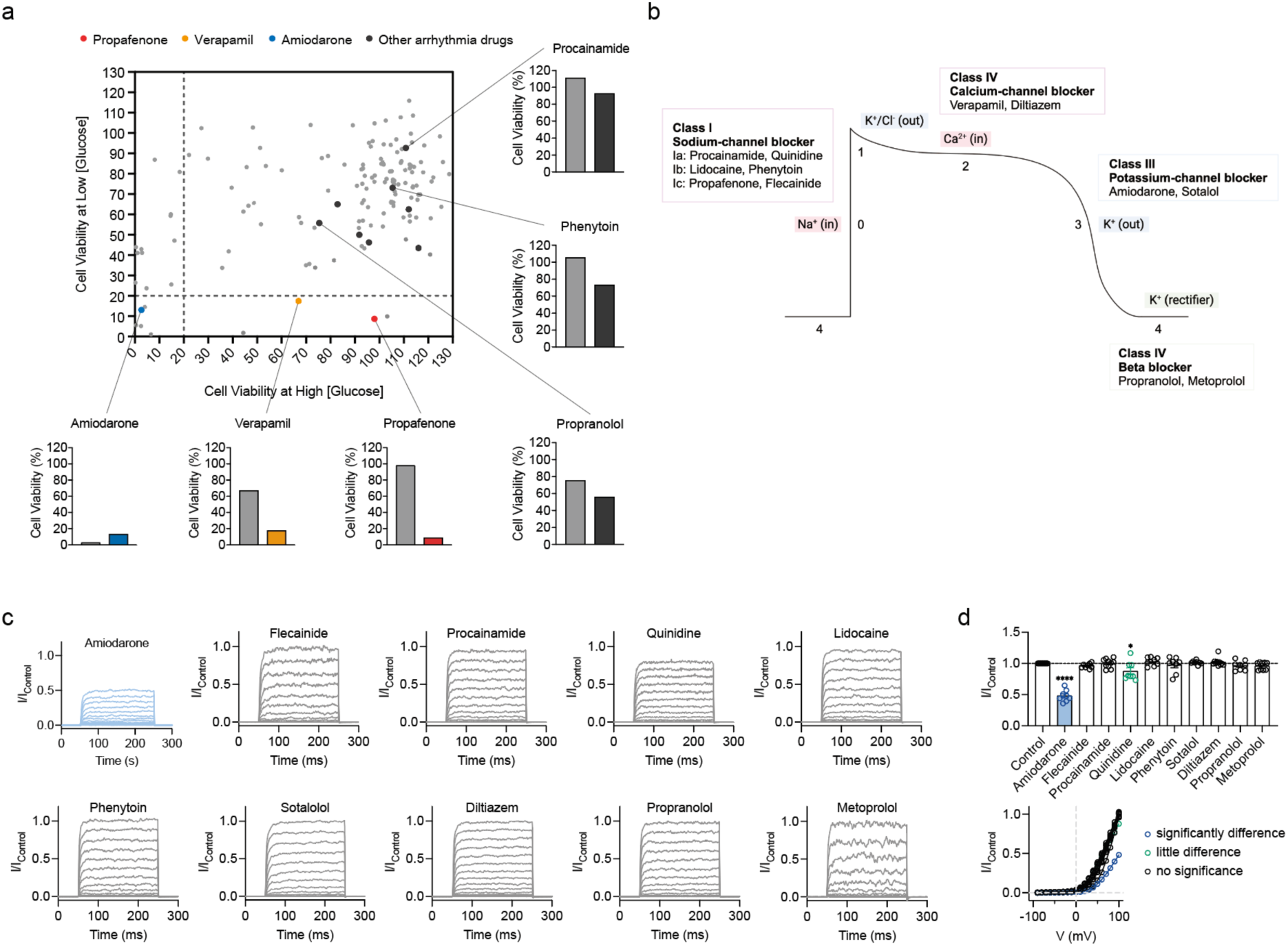
Validation of CM-SLP using other anti-arrhythmic drugs included in CM-SLP. **a** Dot plot of FDA-approved oncology drugs from the CM-SLP^glu^ screen showing relative cell viability under high glucose (x-axis) versus low glucose (y-axis). Procainamide, quinidine, lidocaine, phenytoin, sotalol, diltiazem, propranolol, metoprolol (quadrant 1, black), amiodarone (quadrant 3, blue), propafenone (quadrant 4, red), and verapamil (quadrant 4, orange) are marked. **b** Schematic graph of anti-arrhythmic drugs affecting cardiac action potentials. **c** Representative traces of normalized whole-cell K^+^ currents in the presence of the indicated drugs. Intracellular Ca^2+^ was fixed at 1 μM, and all drugs were added to the extracellular side. Drugs were used at the following concentrations: flecainide, 20 μM; procainamide, 20 μM; quinidine, 20 μM; lidocaine, 20 μM; phenytoin, 20 μM; sotalol, 20 μM; diltiazem, 20 μM; propranolol, 20 μM; and metoprolol, 20 μM. **d** Normalized currents and voltage-current relationships for whole-cell K^+^ currents in the presence of the indicated drugs. Normalized currents at 100 mV in the presence of amiodarone, 0.481 ± 0.033 (n = 8); flecainide, 0.962 ± 0.014 (n = 8); procainamide, 1.002 ± 0.023 (n = 10); quinidine, 0.877 ± 0.053 (n = 8); lidocaine, 1.030 ± 0.024 (n = 8); phenytoin, 0.976 ± 0.045 (n = 8); sotalol, 1.015 ± 0.011 (n = 8); diltiazem, 1.019 ± 0.021 (n = 10); propranolol, 0.966 ± 0.027 (n = 8); and metoprolol, 0.960 ± 0.020 (n = 9). In c and d, the currents were evoked by −90 mV pre-pulse for 50 ms, then −90 to 100 mV test pulses for 200 ms with 10-mV increments, and then −90 mV post-pulse for 50 ms. The channel activity was calculated as the average current density during the last 50 ms of each test pulse.

**Supplementary Fig. 5:**
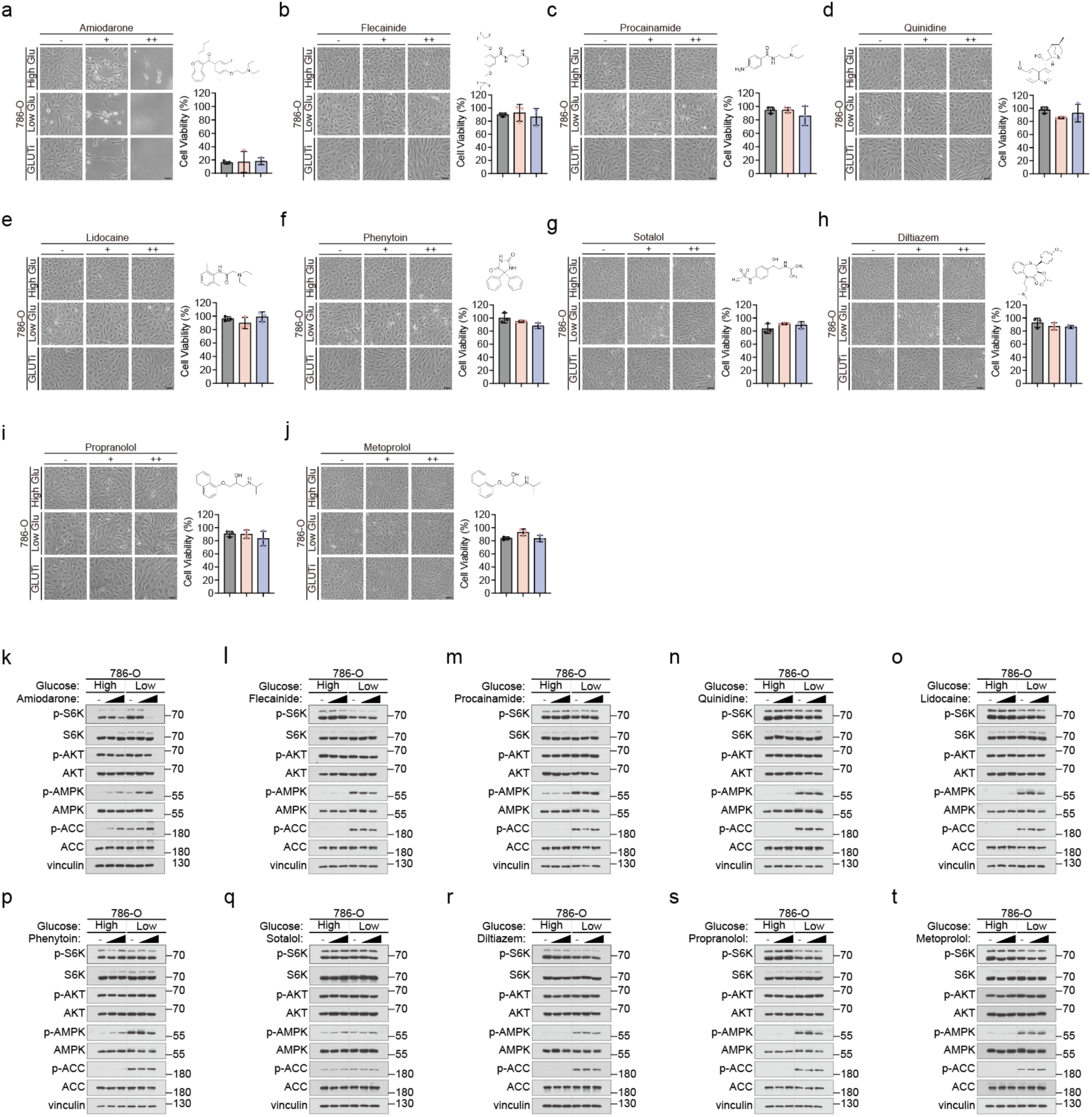
Anti-arrhythmic drugs fail to exhibit synergistic effects in the absence of K_ca_ channels. **a** Viability of 786-O cells subjected to amiodarone (0, 10, 30 μM, 48 h) treatment in the presence of high glucose (25 mM), low glucose (1 mM), or the GLUT inhibitor BAY-876 (3 μM). **b** Viability of 786-O cells subjected to flecainide (0, 10, 30 μM, 48 h) treatment in the presence of high glucose (25 mM), low glucose (1 mM), or the GLUT inhibitor BAY-876 (3 μM). **c** Viability of 786-O cells subjected to procainamide (0, 10, 30 μM, 48 h) treatment in the presence of high glucose (25 mM), low glucose (1 mM), or the GLUT inhibitor BAY-876 (3 μM). **d** Viability of 786-O cells subjected to quinidine (0, 10, 30 μM, 48 h) treatment in the presence of high glucose (25 mM), low glucose (1 mM), or the GLUT inhibitor BAY-876 (3 μM). **e** Viability of 786-O cells subjected to lidocaine (0, 10, 30 μM, 48 h) treatment in the presence of high glucose (25 mM), low glucose (1 mM), or the GLUT inhibitor BAY-876 (3 μM). **f** Viability of 786-O cells subjected to phenytoin (0, 10, 30 μM, 48 h) treatment in the presence of high glucose (25 mM), low glucose (1 mM), or the GLUT inhibitor BAY-876 (3 μM). **g** Viability of 786-O cells subjected to sotalol (0, 10, 30 μM, 48 h) treatment in the presence of high glucose (25 mM), low glucose (1 mM), or the GLUT inhibitor BAY-876 (3 μM). **h** Viability of 786-O cells subjected to diltiazem (0, 10, 30 μM, 48 h) treatment in the presence of high glucose (25 mM), low glucose (1 mM), or the GLUT inhibitor BAY-876 (3 μM). **i** Viability of 786-O cells subjected to propranolol (0, 10, 30 μM, 48 h) treatment in the presence of high glucose (25 mM), low glucose (1 mM), or the GLUT inhibitor BAY-876 (3 μM). **j** Viability of 786-O cells subjected to metoprolol (0, 10, 30 μM, 48 h) treatment in the presence of high glucose (25 mM), low glucose (1 mM), or the GLUT inhibitor BAY-876 (3 μM). **k-t** Immunoblotting analysis of cancer-related signaling pathways in cells treated with the indicated drugs (0, 10, 30 μM, 8 h) in the presence of either high (25 mM) or low (1 mM) glucose. -, negative control; low dose, 10 μM; high dose, 30 μM.

**Supplementary Fig. 6:**
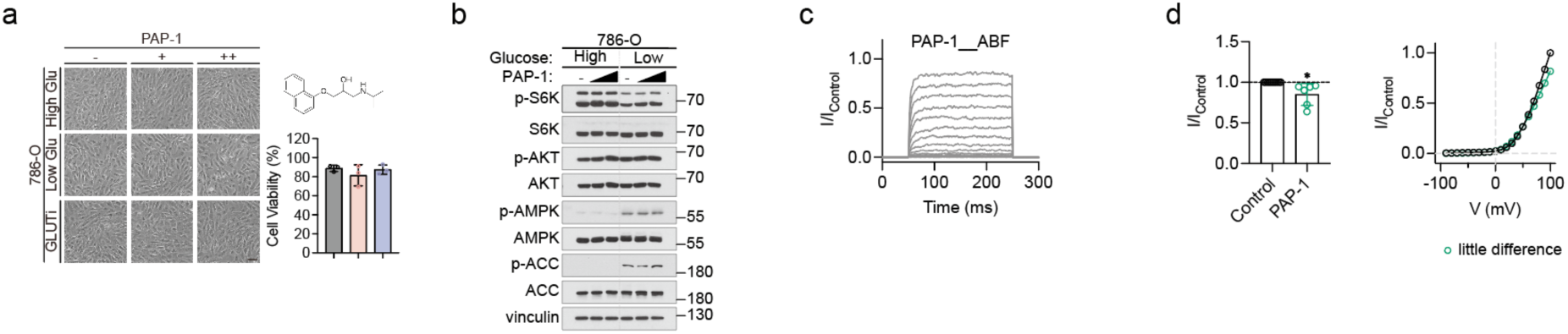
The voltage-gated potassium channel inhibitor PAP-1, which lacks anticancer efficacy, does not target the mTOR pathway or K_ca_ channels. **a** Viability of 786-O cells subjected to PAP-1 (0, 10, 30 μM, 48 h) treatment in the presence of high glucose (25 mM), low glucose (1 mM), or the GLUT inhibitor BAY-876 (3 μM). **b** Immunoblotting analysis of cancer-related signaling pathways in cells treated with PAP-1 (0, 10 30 μM, 8 h) combined with either high (25 mM) or low (1 mM) glucose. -, negative control; low dose, 10 μM; high dose, 30 μM. **c** Representative traces of normalized whole-cell K^+^ currents in the presence of PAP-1. Intracellular Ca^2+^ was fixed at 1 μM, and the drugs were added to the extracellular side. PAP-1 concentrations; 60 μM. **d** Normalized currents and voltage-current relationships for whole-cell K^+^ currents with chemicals. Normalized current at 100 mV; PAP-1, 0.852 ± 0.055 (n = 6). In c and d, currents were evoked by −90 mV pre-pulse for 50 ms, then −90 to 100 mV test pulses for 200 ms with 10-mV increments, and then −90 mV post-pulse for 50 ms. Channel activity was calculated as the average current density during the last 50 ms of each test pulse.

**Supplementary Figure 7.**
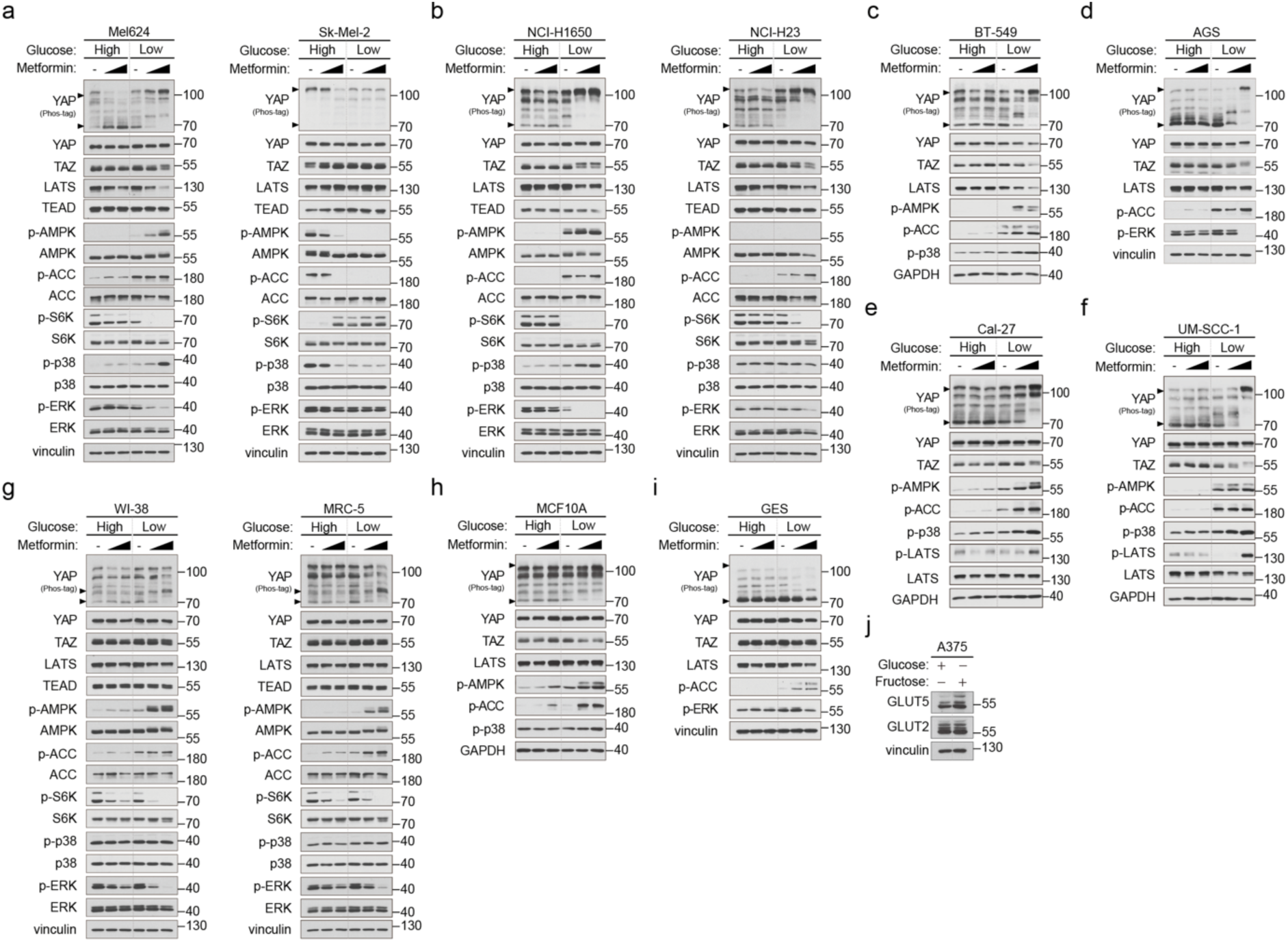
Additional cancer types that show perturbed signaling pathways in response to the biguanide/low glucose combination. **a** Immunoblotting analysis of melanoma cells treated with metformin (0, 3, 10 mM, 8 h) in high (25 mM) or low (1 mM, 11 h) concentrations of glucose. **b** Immunoblotting analysis of lung cancer cells treated with metformin (0, 3, 10 mM, 8 h) in high (25 mM) or low (1 mM, 11 h) concentrations of glucose. **c** Immunoblotting analysis of normal lung cells treated with metformin (0, 3, 10 mM, 8 h) in high (25 mM) or low (1 mM, 11 h) concentrations of glucose. **d** Immunoblotting analysis of breast cancer cells treated with metformin (0, 3, 10 mM, 8 h) in high (25 mM) or low (1 mM, 11 h) concentrations of glucose. **e** Immunoblotting analysis of non-cancer breast cells treated with metformin (0, 3, 10 mM, 8 h) in high (25 mM) or low (1 mM, 11 h) concentrations of glucose. **f** Immunoblotting analysis of gastric cancer cells treated with metformin (0, 3, 10 mM, 8 h) in high (25 mM) or low (1 mM, 11 h) concentrations of glucose. **g** Immunoblotting analysis of normal gastric cells treated with metformin (0, 3, 10 mM, 8 h) in high (25 mM) or low (1 mM, 11 h) concentrations of glucose. **h** Immunoblotting analysis of oral cancer cells treated with metformin (0, 3, 10 mM, 8 h) in high (25 mM) or low (1 mM, 11 h) concentrations of glucose. **i** Immunoblotting analysis of head and neck cancer cells treated with metformin (0, 3, 10 mM, 8 h) in high (25 mM) or low (1 mM, 11 h) concentrations of glucose. **j** Immunoblotting analysis of A375 treatment in the presence of high glucose (25 mM) or high fructose (50 mM).

**Supplementary Fig. 8:**
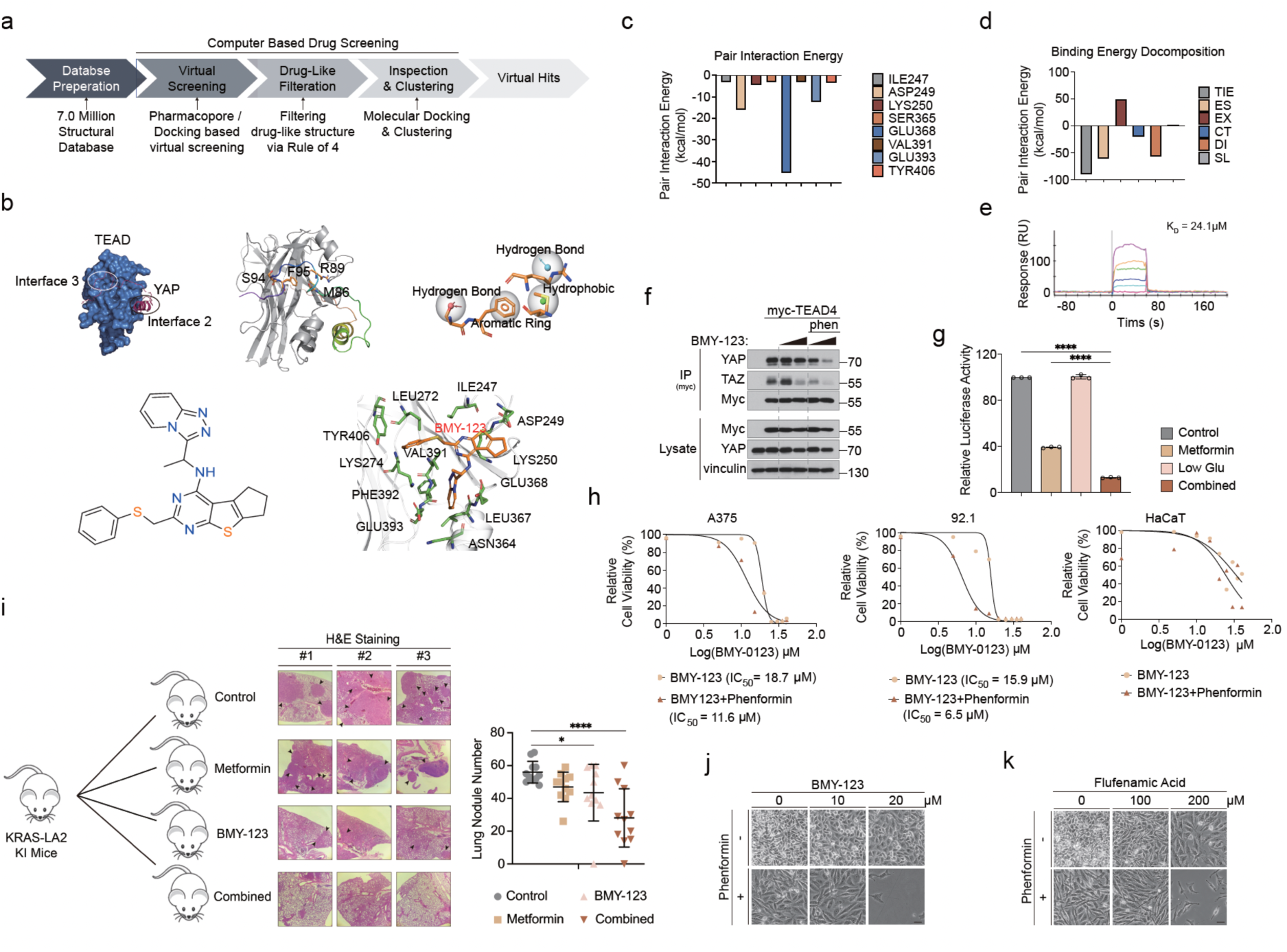
Redirecting metabolic interventions toward TEAD targeted therapy improves the clinical applicability of CM-SLP. **a** Schematic overview of the virtual drug screen used to identify TEAD inhibitors. **b** Schematic overview of the YAP-TEAD binding interfaces (blue, TEAD1; magenta, YAP1, PDB code: 3KYS) and the detailed pharmacophore structure of interface 3 used for virtual screening (top row). Chemical structure of BMY-123 and the residues in TEAD1 interface 3 that interact with BMY-123 are shown (bottom row). **c** Measurements of pair interaction energy (PIE) for the significant residues in TEAD1 interface 3. **d** FMO results for the interaction between TEAD1 interface 3 and BMY-123. Measurements of binding energy decomposition. The total interaction energy (TIE), electrostatic (ES), exchange repulsion (EX), charge transfer (CT), dispersion (DI), and solvation energy terms (SL) are shown. **e** **f** Immunoprecipitation analysis of phenformin/BMY-123-induced disruption of the YAP/TAZ-TEAD interaction in HEK293A cells. Phenformin, 10 μM; BMY-123, 10 μM, 30 μM. **g** TEAD luciferase reporter activity from the 8xGTIIC plasmid expressed in HEK293A cells subjected to phenformin/BMY-123. Y-axis, relative luciferase units (RLU). n = 3. ****, p < 0.0001. Statistics were analyzed via one-way ANOVA with Bonferroni corrections for multiple comparisons. **h** Measurements of cell viability and IC_50_ made via MTT assays on cells subjected to BMY-123 alone or the combination of phenformin/BMY-123. Phenformin, 100 μM. **i** Schematic showing K-Ras^LA2^ lung cancer model mice given intraperitoneal injections of metformin/BMY-123 (left). H&E staining of mouse lung sections (middle). Arrows indicate lung tumor nodules. Counting lung tumor nodules in each group (right). Metformin, 250 mg/kg/day; BMY-123, 10 mg/kg/day *, p < 0.05; ****, p < 0.001. Statistics were analyzed via one-way ANOVA with Bonferroni corrections for multiple comparisons. **j** Synergetic cytotoxicity of phenformin with BMY-123. Phenformin was administered at 100 μM with BMY-123 (0, 10, 20 μM) for 20 h. **k** Synergetic cytotoxicity of phenformin with BMY-123. Phenformin was administered at 100 μM with flufenamic acid (0, 100, 200 μM) for 20 h.

## Methods

### CM-SLP high-throughput screening

A375 and HaCaT cells were plated on 96-well plates at a density of 1 x 10^3^ cells per well. After 24 h incubation, the plates were washed 3 times with PBS (200 μl/well), and the cell culture medium was replaced with either high or low nutrient-containing medium. For the CM-SLP^glu^ platform, we used media containing 25 mM and 1 mM glucose; for the CM-SLP^gln^ platform, we used 2 mM and 0 mM glutamine; and for the CM-SLP^FA^ platform, we used normal versus charcoal-filtered fatty acid-depleted serum. The cells were then treated with an approved drug library (Targetmol, Boston, MA) containing 1,813 compounds, each at 100 μM. For each platform, cells were treated with drugs for 36 h (CM-SLP^glu^), 18 h (CM-SLP^gln^), and 60 h (CM-SLP^FA^), then cell viability was measured with CellTiter 96*®* AQueous One Solution Assay (Promega, Madison, WI), according to manufacturer’s instructions. Absorbance was then measured at 490nm using Infinite 200 PRO plate reader (Tecan Group Ltd., Mannedorf, Switzerland). CM-SLP scores were calculated by following equation: CM-SLP score = log_0.5_([cell viability at high nutrient]/[cell viability at low nutrient]).

### Cell Culture

All cell lines were maintained at 37°C under 5% CO_2_. The A375, HaCaT, HEK293A, MEF, and Mel624 cell lines were cultured in DMEM (Cytiva, SH30022.01) containing 10% FBS (Hyclone, SV30207.02) and 50 μg/ml penicillin/streptomycin (Gibco, 15140122). 786-O, NCI-H1975, NCI-H358, BEAS-2B, BT549, SNU885, SNU878, A549, NCI-H1650, NCI-H23, 92.1, y-Meso-26B, K562, Molm14, Malm6, THP-1, NCI-H69, and NCI-H854 cell lines were cultured in RPMI (Hyclone, SH30027.01) containing 10% FBS (Hyclone, SV30207.02) and 50 μg/ml penicillin/streptomycin (Gibco, 15140122). The HEMn and HEMa cell lines were cultured in Dermal Cell Basal Medium (ATCC, PCS-200-030) complemented with Melanocyte Growth Kit (ATCC, PCS-200-041) and Adult Melanocyte Growth Kit (ATCC, PCS-200-042), respectively. The HREC cell line cultured in Renal Epithelial Cell Basal Medium (ATCC, PCS-400-030) complemented with Renal Epithelial Cell Growth Kit (ATCC, PCS-400-040). MCF10A cell line was cultured in DMEM-F12 (Hyclone, SH30023.01) supplemented with 5% horse serum (Gibco, 16050122), 20 ng/ml EGF (Peprotech, 315-09), 0.5 μg / ml hydrocortisone (Sigma, H0135), 100 ng/ml cholera toxin (Sigma, SAE0069) and 10 μg/ml insulin (Sigma, I9278). Sk-mel-2 and MRC-5 cell lines were cultured in MEM (Cytiva, SH30024,01) containing 10% FBS (Hyclone, SV30207.02) and 50 μg/ml penicillin/streptomycin (Gibco, 15140122). No cell lines used in this study were found in the database of commonly misidentified cell lines maintained by ICLAC and NCBI Biosample. To induce metabolic stress, cells were starved of glucose, glutamine, and fatty acids by switching them to glucose-free DMEM (Gibco, 1196025), glutamine-free DMEM (Hyclone, SH30081.01), and charcoal-stripped FBS (Thermo, A3382101), respectively.

### CRISPR KO cell lines

GPI, AMPKα1/2, LATS1/2, and LKB1 knock-out cell lines were engineered using the CRISPR-Cas9 gene-editing system. The guide RNA (gRNA) sequences targeting these genes were designed using the CRISPR design tool available at http://www.rgenome.net/cas-designer/. The respective gRNA sequences employed were as follows: for GPI #1 5’-GGTTCTTGGAGTAATCCACC-3’, #2 5’-GCATCACGTCCTCCGTCACC-3’; for AMPKα1 #1 5’-GCGAGCTTCGTCCTCATGCAGGG-3’, #2 5’-TACTCAATCGACAGAAGATT-3’; for AMPKα2 #1 5’-GAAGATCGGACACTACGTGC-3’, #2 5’-CTACGTGCTGGGCGACACGC-3’; for LATS1 #1 5’-GCAGCCATCTGCTCTCGTCG-3’, #2 5’-GGAGGTGGAGTTGTACCTCT-3’; for LATS2 #1 5’-GTAGGACGCAAACGAATCGC-3’, #2 5’-TAGCCCCTGAACCGAAGACT-3’; and for LKB1 #1 5’-AGCTTGGCCCGCTTGCGGCG-3’, #2 5’-CCACCGCATCGACTCCACCG-3’. The custom-designed gRNA sequences targeting GPI, AMPKα1, AMPKα2, LATS1, LATS2, and LKB1 were cloned into the px459 vector (Addgene plasmid No. 62988). Following DNA sequencing to verify the accuracy of the insertions, the plasmids containing the desired sequences were transferred to DH5α *Escherichia coli* to complete cloning. The cloned px459 vectors were then transfected into HEK293A cells. Post-transfection, the cells were subjected to puromycin selection. Individual HEK293A clones were separated into 96-well plates using a BD FACS Aria™ III cell sorter (BD Biosciences, San Jose, CA, USA). The successful generation of knock-out clones was subsequently confirmed through western blot analysis. p53 knock-out and p53/TSC2 double knock-out MEF cell lines were a gift from Young Jin Jang (Major of Food Science & Technology, Seoul Women’s University, Seoul 01797, Republic of Korea).

### Chemical agents

Propafenone hydrochloride (P4670), metformin hydrochloride (#PHR1084), warfarin (#A2250), fenbendazole (#PHR1832), phenformin hydrochloride (#PHR1573), fructose (#F3510), oligomycin A (#75351) and BAY-876(SML1774) were purchased from Sigma Aldrich. Verapamil hydrochloride (#14288), paxilline (#11354), flecainide acetate (#20388), procainamide hydrochloride (#24359), quinidine (#20356), lidocaine (#20081), phenytoin (#24037), sotalol hydrochloride (#16136), diltiazem hydrochloride (#20079), nilutamide (#23953), cisapride (#21657), dabrafenib (#16989), trametinib (#16292), and valdecoxib (#10006120) were purchased from Caymanchem. Tram-34 (S1160), tetraethylammonium chloride (S4489), propranolol hydrochloride (S4076), metoprolol (S5430), and temsirolimus (S1044) were purchased from Selleckchem. Rapalink-1 (A8764) was purchased from Biotrend.

### Immunofluorescence microscopy

Cells were seeded onto 12-well plates on coverslips 1 day prior to experimentation. Coverslips were pretreated with poly-L-ornithine solution (Sigma, P4957) diluted 1:20 at 37 °C for 15 min, followed by a quick phosphate buffer saline (PBS) wash prior to cell seeding. The cells were then fixed in 4% paraformaldehyde (Thermo, 28908) for 20 min and permeabilized in 0.1% Triton-X/PBS for 5 min. The cells were blocked in 3% BSA/PBS for 30 minutes and incubated overnight at 4 °C in primary antibodies diluted in 3% BSA/PBS. Then the cells were incubated for 2 h in secondary antibodies diluted in 3% BSA/PBS. The slides were mounted in Prolong gold antifade reagent with DAPI (Invitrogen, P36930). Single Z section images at the same cellular level were captured with a confocal microscope (Leica Microsystems, SP8). The images depicted in the figures were processed and exported using ImageJ.

### Immunoprecipitation and immunoblotting

Immunoblotting was performed using a standard protocol. Phos-tag reagents were purchased from Wako Chemicals, and gels containing phos-tag were prepared according to the manufacturer’s instructions. For immunoprecipitations, cells were rinsed twice with ice-cold PBS and lysed in ice-cold lysis buffer (0.15 M NaCl, 0.05 M Tris-HCl, 0.5% Triton X-100, and one tablet each of EDTA-free protease and phosphatase inhibitors (Thermo, 78446). For immunoprecipitations, anti-Myc magnetic beads (Thermo, 88842) were added to the lysates and incubated with rotation overnight at 4 °C. The resulting immunoprecipitates were washed three times with lysis buffer. The immunoprecipitated proteins were then denatured with the addition of sample buffer and subjected to boiling for 7 min. Then, they were resolved via 8% SDS-PAGE and analyzed via western blot.

### Antibodies

The following antibodies were purchased from Cell Signaling and used at the indicated dilutions for western blot analysis, immunohistochemistry, and immunofluorescence: p-S6K (#9234), S6K (#2708), p-AKT (#4060), AKT (#4685), p-AMPK (#2535), AMPK (#2603), p-ACC (#11818), ACC (#3676), YAP (#14074), TAZ (#4883), LATS (#9153), pan-TEAD (#13295), p-p38 MAPK (#4511), p38 MAPK (#8690), P-ERK 1/2 (#4377), ERK 1/2 (#4695), p-FAK (#3283), FAK (#3285), p-LATS (#8654), LATS (#3477), IgG (#2729). The following antibodies was purchased from Santa Cruz Biotechnology and used at the indicated dilutions for 853 western blot analysis and immunofluorescence: Myc-HRP (sc-40), GAPDH (#sc-25778), Vinculin (#sc-7364). Paxillin (#610569) antibody was purchased from BD bioscience.

Rhodamin-Phalloidin (#R415) was purchased from ThermoFisher. The following antibodies were purchased from Abcam and used at the indicated dilutions for western blot analysis: GLUT5 (ab279363) and GLUT2 (ab192599).

### Extracellular flux analysis

786-O renal cancer cells (3 x 10^3^) were distributed into the individual wells of 96-well plates (Agilent Technologies, 103774-100), with six wells designated for each experimental group. To serve as a negative control, the four corners of the plate were deliberately left empty of cells. In these regions, only Seahorse media was provided. After 24 hours of incubation, each well was washed with 100 μL of phosphate-buffered saline (PBS) and then resuspended in prewarmed (37°C) DMEM media with either 25 mM glucose or 0.5 mM glucose, 40 μM propafenone or 30 μM paxilline, and DMSO for each control group. After 24 hours of pre-treatment, the 786-O cells were washed with PBS and then resuspended in Seahorse Basal Medium supplemented with 2 mM glutamine and either 25 mM glucose or 0.5 mM glucose at pH 7.4 for ATP Real-Time Assays.

Using a Seahorse XF96 Analyzer (Agilent Technologies, USA), we measured extracellular acidification rate (ECAR) and oxygen consumption rate (OCR) in accordance with the manufacturer’s instructions. Baseline OCR and ECAR values were obtained through three initial measurements, followed by the injection of Seahorse XF Real-Time ATP Rate Assay Kit reagents. The final concentrations of these reagents (1.5 μM Oligomycin-A, 0.5 μM Rotenone, and 0.5 μM Antimycin A) were achieved using the Seahorse XF Real-Time ATP Rate Assay Kit (#103592-100) from Agilent. Subsequent measurements were facilitated by the ATP Real-Time Rate Assay Generator (Agilent Technologies). Afterward, 786-O cells were normalized to the protein concentration per well, as measured via a Bradford assay kit (#5000006) purchased from Bio-rad.

The real-time analysis of ATP production was conducted using the Seahorse XFp Analyzer (Agilent Technologies, USA) and the Seahorse XFp real-time ATP rate assay kit (Agilent Technologies, 103591-100) according to the manufacturer’s guidelines. A375 cells were seeded into XFp cell culture miniplates (Agilent Technologies, 103025-100) at a density of 1 x 10^4^ cells per well. After 24 hours of incubation, each well was washed with 100 μL of PBS. Then, the medium was replaced with either 25 mM or 0 mM glucose-containing medium. Three hours after this medium change, 25 mM metformin was added and incubated for 8 hours. After this drug incubation period, each miniplate was placed in the XFp analyzer along with the XFp sensor cartridge that contained the drug from the assay kit to measure real-time ATP production. These analyses were conducted according to the manufacturer’s instructions.

### ATP Assay

ATP levels were measured to determine whether the reduction in ATP production induced by combined treatment with glucose restriction and metformin could be rescued by fructose. A375 cells were seeded into 12-well plates at 5 x 10^5^ cells per well. After a 24-hour incubation, the cells were washed twice with PBS. Then, the medium was replaced with medium containing either 25 mM or 0 mM glucose. Three hours after the medium change, 10 mM metformin was added and incubated for 8 hours. ATP levels were then measured with the ATP Detection Assay Kit (Cayman Chem, 700410) according to the manufacturer’s instructions.

### Cell Viability Assay

Cell viability assays were conducted to determine whether the cell death effect of combined treatment with glucose restriction and metformin is rescuable by fructose. A375 cells were seeded into 12-well plates at 5 x 10^5^ cells per well. After a 24-hour incubation, the cells were washed twice with PBS and then the medium was replaced with medium containing either 25 mM or 0 mM glucose. Three hours after the medium change, 10 mM metformin was added and allowed to incubate for 16 hours. Subsequently, photographs of the cells were taken. After a single PBS wash, the medium was replaced with DMEM containing MTT at a 1:100 dilution from a 12 mM stock solution (Thermo, M6494). Four hours after the medium change, the medium was removed, and the cells were solubilized with 300 μL of DMSO to measure absorbance and analyze cell viability.

### RNA extraction, cDNA synthesis, and quantitative real-time PCR (qRT-PCR)

Cells were harvested for RNA extraction using the RNeasy Plus mini kit (QIAGEN, 74136). RNA samples were reverse transcribed to complementary DNA (cDNA) using iScript reverse transcriptase (Bio-Rad, 1708891). qRT-PCR was performed using the KAPA SYBR FAST qPCR kit (Kapa Biosystems, KK4605) and the StepOnePlus Real-Time PCR System (Applied Biosystems). The qRT-PCR primers were as follows: CTGF – F, 5’-ATGACACTGTTCAGGAATCG - 3’; CTGF – R, 5’-CAAATTCACTTGCCACAAGC; ANKRD1 – F, 5’-CACTTCTAGCCCACCCTGTGA -3’; ANKRD1 – R, 5’-CCACAGGTTCCGTAATGATTT -3’; GAPDH – F, 5’-ACCAGGTGGTCTCCTTCGAC -3’; GAPDH– R, 5’-TGCTGTAGCCAAATTCGT -3’.

### Animal experiments

NOD/SCID mice were purchased from JaBio (South Korea). For the tumor xenograft models, 786-O renal cancer cells (5 x 10^6^) were injected subcutaneously into the right flank of 6-week-old male nude mice. Seven days after the injections, the mice were assigned randomly to treatment groups, eight mice per group. For *in vivo* drug treatments, NOD/SCID mice were given intraperitoneal injections 4 days a week with propafenone (30 mg/kg), temsirolimus (0.3 mg/kg), propafenone (30 mg/kg) with temsirolimus (0.3 mg/kg), Bay-876 (5 mg/kg), propafenone with Bay-876 (5 mg/kg), or vehicle (PBS). Propafenone, temsirolimus, and Bay-876 were diluted in PBS and thoroughly sonicated. Mouse weights were measured every week. Tumor weight and volume were measured after the animals were sacrificed. Mice were sacrificed 6 weeks after the experiment began. Balb/C nude mice were purchased from JaBio (South Korea). For the tumor xenograft models, A375 melanoma cells (2 x 10^5^) and NCI-H1975 lung cancer cells (5 x 10^5^) were injected subcutaneously into the right flank of 6-week-old male nude mice. When the resulting tumors reached 100 mm^2^, the mice were assigned randomly to treatment groups. The investigators were not blind to these allocations during the experiments or the outcome assessments. To manipulate mouse blood glucose, mice in the carbohydrate-free group were fed carbohydrate-free chow according to the average weight of AIN-93M control chow. For *in vivo* metformin treatments, nude mice were given daily intraperitoneal injections of metformin diluted in PBS at doses of 200 mg/kg. Tumor volume and mouse weight were measured every week (volume = width * width * height/2) and blood glucose was measured daily with a blood glucose monitor (Glucodoctor). The mice were euthanized 6 weeks after tumor engraftment. K-Ras-LA2 mice were purchased from the Jackson Laboratory. These mice were bred in a specific pathogen-free (SPF) facility in the Yonsei Laboratory Animal Research Center. In vitro fertilization (IVF) was used to augment the cancer model mice. They were fed a normal chow diet (PicoLab® Rodent Diet 20, Orient Bio, Inc.) and maintained under a 12-hour light-dark cycle in the SPF facility at 23°C and 40–60% humidity. For the *in vivo* BMY-123 and metformin treatments, LA2 mice were given daily intraperitoneal injections with 10 mg/kg of BMY-123 and 200 mg/kg metformin. BMY-123 was diluted in PBS and thoroughly sonicated. Mice were sacrificed 10 weeks after the experiment began. All animal experiments were approved by the Yonsei University Institutional Animal Care and Use Committee (Documentation #201610-435-02).

### Patch Clamp Solutions

Whole-cell voltage clamp experiments were conducted using a basal extracellular solution containing 130 mM NaCl, 4 mM KCl, 10 mM HEPES, 10 mM glucose, 1 mM CaCl_2_, and 1 mM MgCl_2_, with its pH adjusted to 7.4 using NaOH and its osmolarity adjusted to ∼310 mOsM with an appropriate amount of sorbitol. The basal pipette solution with 1 μM free Ca^2+^ contained 130 mM KCl, 10 mM HEPES, 5 mM EGTA, and 4.37 mM CaCl_2_, and 3 mM Mg-ATP, with its pH adjusted to 7.2 using KOH and its osmolarity adjusted to ∼290 mOsM with an appropriate amount of sorbitol. Free Ca^2+^ was calculated using the WEBMAX-C software (C. Patton, Stanford University; https://somapp.ucdmc.ucdavis.edu/pharmacology/bers/maxchelator/webmaxc/webmaxcS.htm). The pipette solution with zero free Ca^2+^ was made by omitting CaCl_2_.

For recording voltage-activated Na^+^ currents, the extracellular solution contained 130 mM NaCl, 4 mM CsCl, 1 mM MgCl_2_, 10 mM glucose, 10 mM HEPES, and 100 μM CdCl2, with its pH adjusted to 7.4 using NaOH and its osmolarity adjusted to ∼310 mOsm with sorbitol. The intracellular solution contained 117 mM CsCl, 20 mM NaCl, 1 mM MgCl_2_, 5 mM HEPES, and 5 mM EGTA, with its pH adjusted to 7.2 using KOH and its osmolarity adjusted to ∼290 mOsm with sorbitol.

For recording voltage-activated Ca^2+^ currents, the extracellular solution contained 120 mM NaCl, 5 mM CsCl, 0.5 mM MgCl_2_, 10 mM BaCl_2_, 10 mM HEPES, 10 mM tetraethylammonium (TEA)- Cl, and 10 mM glucose, with its pH adjusted to 7.4 using NaOH and its osmolarity adjusted to ∼300 mOsM with sorbitol. The intracellular solution contained 100 mM CsOH, 100 mM aspartic acid, 32 mM CsCl, 10 mM EGTA, 10 mM HEPES, and 5 mM Mg-ATP, with its pH adjusted to 7.2 using CsOH and its osmolarity adjusted to ∼290 mOsm using sorbitol.

### Electrophysiology

Cells were transferred to a bath perfused at 5 mL/min and mounted on the stage of an inverted microscope (Nikon, Japan). Microglass pipettes (World Precision Instruments, USA) were fabricated using a PP-830 single-stage glass microelectrode puller (Narishige, Japan), with a resistance of 2–5 MΩ. The liquid junction potential was rectified using an offset circuit prior to each recording. Currents were recorded using an Axopatch 200B amplifier (Molecular Devices, USA) and Digidata 1440A interface (Molecular Devices), digitized at 20 kHz, and low-pass filtered at 2 kHz using pClamp software 10.7 (Molecular Devices). All recordings were performed at room temperature (22–25°C). The whole-cell voltage clamp configuration was verified by measuring the series resistance to <10 MΩ, which was compensated before each recording.

In the whole-cell configuration, K^+^ currents were recorded at a holding potential of –60 mV, and chemicals were applied after >5 min of break-in to reduce the effects of channel activation. The chemicals were tested for >5 min to obtain a stable current level. To measure the channel activity on blocker administration experiments, cells were held at currents were evoked by −90 mV pre-pulse for 50 ms, then −90 to 100 mV test pulses for 200 ms with 10 mV increment, then −90 mV post-pulse for 50 ms. Channel activity was calculated as the average current density in the last 50 ms of each test pulse. All experiments were performed in independent biological replicates, and the mounting chamber was replaced with new cells after each chemical.

### Protein structure preparation

The X-ray crystal structure of human YAP bound to TEAD1 was retrieved from the Protein Data Bank (PDB ID: 3KYS). All the missing side chains were filled in using Prime implemented in Maestro program^55^. Hydrogen atoms were added to the crystal structure at pH 7.0 and their positions were optimized with PROPKA implemented in Maestro program^56^. Then the restrained energy minimizations were performed with an OPLS3 force field set to 0.3 Å root mean square deviation^57^.

### Virtual screening

To find compounds that inhibit the protein-protein interaction (PPI) between TEAD1 and YAP, pharmacophore-based virtual screening was performed on our in-house synthetic database using Phase implemented in the Schrödinger suite^58,59^. Pharmacophores were generated for the PPI between TEAD1 and YAP. The top-hit inhibitor, BMY-123, was selected from the pharmacophore-based screening. Then molecular docking performed using Glide implemented in the Schrödinger suite^60^. The docking position of BMY-123 bound to TEAD1 was selected using its glide score, e-model score, and by visual inspection.

### Fragment molecular orbitals

To analyze the interactions between TEAD1 and BMY-123 at the molecular level, we performed an ab initio fragment molecular orbital (FMO) analysis. All FMO calculations were performed using GAMESS^61^. The energy minimization of the top-hit docking pose of BMY123 was performed at FMO-DFTB3/D/PCM level with the third order corrected density functional tight-binding (DFTB3) method using the 3OB parameter set^62,63^, UFF-type dispersion correction (D)^64^, and polarizable continuum model (PCM)^65^. In energy minimization, the residues within 10.4 Å from the ligand were 908 included and fixed, and only the ligand was allowed to remain fully flexible. The energy minimization calculations converged in 61 steps.

An energy decomposition analysis was performed with the energy minimized structure at FMO-MP2/PCM level with the second order Møller-Plesset perturbation theory (MP2)^66^, and a polarizable continuum model (PCM)^67^ using the 6-31G** basis set. All the residues in the crystal structure were included in the energy decomposition analysis. The binding affinity of the protein-ligand interaction was approximated as the sum of their pair interaction energies (PIE). Each PIE provided physical details of the protein-ligand interactions^68,69^. The PIEs between fragments in the FMO calculations were decomposed by five energy terms defined by Eq (1): electrostatic (Δ*E^es^*), exchange-repulsion (Δ*E^ex^*), charge transfer with a higher order mixed term (Δ*E^ct+mix^*, dispersion (Δ*E^di^*), and solvation energy (Δ*G_sol_*) from the polarizable continuum model (PCM).

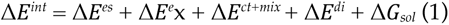

The electrostatic term is mainly derived from the Coulomb interaction between the polarized charge distributions of the fragments. The charge transfer term is derived from the interaction between the occupied orbitals of a donor and the unoccupied orbitals of an acceptor. The electrostatic and charge transfer terms are important for assigning hydrogen bond, salt-bridge, and polar interaction positions. The exchange-repulsion term is derived from the interactions between fragments located in proximity with one another. The dispersion term derived from the interaction of the induced dipole moments of two fragments. The exchange-repulsion term describes steric repulsion, while the dispersion term represents hydrophobic in nature.

### Statistics and reproducibility

All quantitative data were obtained from at least three independent biological replicates. All data are presented as means ± standard deviation (s.d.) unless otherwise noted in the figure legends. Statistical differences between two groups were examined using two-tailed, unpaired Student’s t-tests or one-way analyses of variance (ANOVA) with Bonferroni corrections for multiple comparisons. Statistical tests were performed using the GraphPad Prism 9.0 software (GraphPad Software, CA, USA). Two-sided p-values of less than 0.05 were considered significant. No statistical methods were used to predetermine sample size. Sample size was based on previous experience with experimental variability. Blinding was performed wherever possible during all sample analyses by coding sample identity during data collection and having the following analyses performed by observers without knowledge of or access to the experimental conditions.

## Acknowledgments

This work was supported by grants from the National Research Foundation of Korea (2020M3F7A1094077, 2020M3F7A1094089, 2021R1A2C1010828, RS-2024-00509461 to HWP, 2018R1D1A1B07043856 to HSJ), by a Global Learning & Academic Research Institution for Masters/PhD students and Postdocs (LAMP) Program grant from the National Research Foundation of Korea funded by the Ministry of Education (RS-2024-00442483 to H.W.P.), and by a Brain Korea 21 FOUR Program grant (to S.Y.P., D.H.K., H.-R.K, S.C.C., J.E.P).

## Author contributions

H.W.P., W.Y.P, and J.H.P. designed the experiments. W.N. and D.J. performed CM-SLP drug screen.

J.W.R. performed the patch clamps experiments. K.T.N., J.K., and H.L. performed TEAD inhibitor development. H.-W.L. and S.Y.P. performed the mouse experiments. H.-S.J., K.-L.G., S.-Y.K., and J.E.S. provided resources and performed specific experiments. H.W.P., W.Y.P., and J.H.P. wrote the manuscript

